# WAPL maintains dynamic cohesin to preserve lineage specific distal gene regulation

**DOI:** 10.1101/731141

**Authors:** Ning Qing Liu, Michela Maresca, Teun van den Brand, Luca Braccioli, Marijne M.G.A. Schijns, Hans Teunissen, Benoit G. Bruneau, Elphège P. Nora, Elzo de Wit

**Affiliations:** Division Gene Regulation, Oncode Institute, Netherlands Cancer Institute, Amsterdam, The Netherlands; Gladstone Institutes, San Francisco, USA; Cardiovascular Research Institute, University of California, San Francisco; Department of Pediatrics, University of California, San Francisco

## Abstract

The cohesin complex plays essential roles in sister chromatin cohesin, chromosome organization and gene expression. The role of cohesin in gene regulation is incompletely understood. Here, we report that the cohesin release factor WAPL is crucial for maintaining a pool of dynamic cohesin bound to regions that are associated with lineage specific genes in mouse embryonic stem cells. These regulatory regions are enriched for active enhancer marks and transcription factor binding sites, but largely devoid of CTCF binding sites. Stabilization of cohesin, which leads to a loss of dynamic cohesin from these regions, does not affect transcription factor binding or active enhancer marks, but does result in changes in promoter-enhancer interactions and downregulation of genes. Acute cohesin depletion can phenocopy the effect of WAPL depletion, showing that cohesin plays a crucial role in maintaining expression of lineage specific genes. The binding of dynamic cohesin to chromatin is dependent on the pluripotency transcription factor OCT4, but not NANOG. Finally, dynamic cohesin binding sites are also found in differentiated cells, suggesting that they represent a general regulatory principle. We propose that cohesin dynamically binding to regulatory sites creates a favorable spatial environment in which promoters and enhancers can communicate to ensure proper gene expression.

**HIGHLIGHTS:**

1. The cohesin release factor WAPL is crucial for maintaining a pluripotency-specific phenotype.
2. Dynamic cohesin is enriched at lineage specific loci and overlaps with binding sites of pluripotency transcription factors.
3. Expression of lineage specific genes is maintained by dynamic cohesin binding through the formation of promoter-enhancer associated self-interaction domains.
4. CTCF-independent cohesin binding to chromatin is controlled by the pioneer factor OCT4.

## INTRODUCTION

The ring-shaped cohesin complex is essential for maintaining chromosome organization at the sub-megabase scale. Cohesin is a multimeric complex consisting of SMC1A, SMC3, RAD21 and one SA subunit (SA1 or SA2). In vertebrate genomes, stable chromatin loops are formed between two convergent CTCF binding sites that block cohesin (Rao et al., 2014; Sanborn et al., 2015; de Wit et al., 2015). We and others have recently shown that the 3D genome can be massively re-organized by knocking out or rapidly depleting cohesin subunits, regulators of cohesin or CTCF (Haarhuis et al., 2017; Nora et al., 2017; Rao et al., 2017; Schwarzer et al., 2017; Wutz et al., 2017). Despite severe changes in loop and TAD structure, the effects of 3D genome changes on transcription are either mild or difficult to explain genome-wide (Hyle et al., 2019; Nora et al., 2017; Rao et al., 2017). Despite the existence of specific examples where CTCF assists in bringing promoters and enhancers together to activate gene expression (Hadjur et al., 2009; Paliou et al., 2019), these results cannot be generalized. Although, the role of architectural proteins in genome organization is becoming clearer, the detailed molecular mechanisms of how these proteins contribute to gene regulation is still poorly understood.

This is also the case for the cohesin release factor WAPL, which dissociates cohesin rings from chromatin by creating a DNA exit gate (Chan et al., 2012; Huis in ‘t Veld et al., 2014) and is thereby important for controlling cohesin levels on chromosomes (Kueng et al., 2006). The WAPL protein is required for various cellular process including sister chromatid resolution (Nishiyama et al., 2010) and DNA repair (Misulovin et al., 2018). The cohesin removal function of WAPL is also important in maintaining genome architecture in mammalian cells. Loss of the WAPL protein results in a genome-wide stabilization of cohesin on chromatin, resulting in the formation of vermicelli chromosomes. This state is characterized by increased chromatin loop size, decreased intra-TAD contact frequency and a suppression of compartments (Haarhuis et al., 2017; Wutz et al., 2017). However, also here it remains to be solved how these changes in 3D genome organization affect transcription regulation.

A significant fraction of chromatin-bound cohesin is not bound at CTCF sites, but co-localizes with lineage specific transcription factors and active chromatin features (enhancers) in specific regions of the genome (Faure et al., 2012; Kagey et al., 2010; Nitzsche et al., 2011) and are frequently associated with cell identity genes. The SA2 subunit defines a subset of cohesin complexes that preferably bind to enhancers sequences (Cuadrado et al., 2019; Kojic et al., 2018). CTCF binding sites, on the other hand, seem to be occupied by both SA1 and SA2 containing cohesin. Clearly, different subsets of cohesin are bound to chromatin, which may affect genome function in different ways.

In this study, we employed acute protein depletion to rapidly deplete WAPL in mouse embryonic stem cells (mESCs), enabling us to examine the immediate effects of changes in cohesin binding and 3D genome changes. We identified regions that lose cohesin binding and local chromatin interactions upon WAPL depletion. These regions are frequently located at or adjacent to pluripotency genes and are enriched for pluripotency transcription factor binding sites. Binding of cohesin to pluripotency transcription factor binding sites is dependent on OCT4, but not NANOG. Finally, we show that WAPL dependent cohesin binding sites exist in differentiated cells as well, indicating the general importance of WAPL for transcription regulation in the mammalian genome.

## RESULTS

### WAPL is required for maintaining the pluripotent transcriptional state

We have previously shown the importance of the cohesin release factor WAPL in maintaining physiological 3D genome organization. In order to study the immediate effects of WAPL loss, cohesin stabilization and 3D genome changes on gene expression we created an acute depletion line for WAPL in mESCs. We fused an AID-eGFP sequence at the C-terminus of the endogenous WAPL protein with CRISPR-Cas9 genome editing (Figure 1A and S1A) (Natsume et al., 2016) into a *OsTir1* parental line (Nora et al., 2017). As expected the tagged WAPL protein showed rapid degradation when indole-3-acetic acid (IAA) was added in the culture medium (Figure 1B and 1C). Upon WAPL depletion, we stained for chromatin-bound cohesin subunit RAD21 (also known as SCC1) and observed the formation of the characteristic Vermicelli chromosomes (Tedeschi et al., 2013) (Figure 1C). Nearly complete WAPL depletion was achieved after 45 minutes of IAA treatment (Figure 1D). We performed calibrated ChIP-seq analysis for WAPL and CTCF and found that acute depletion leads to a genome-wide loss of WAPL binding, but has almost no impact on the genome-wide distribution of CTCF (Figure 1E, S1B). Taken together, these results show that our WAPL-AID cell line enables us to study the effects of rapid cohesin stabilization on cellular functions.

**Figure 1:**
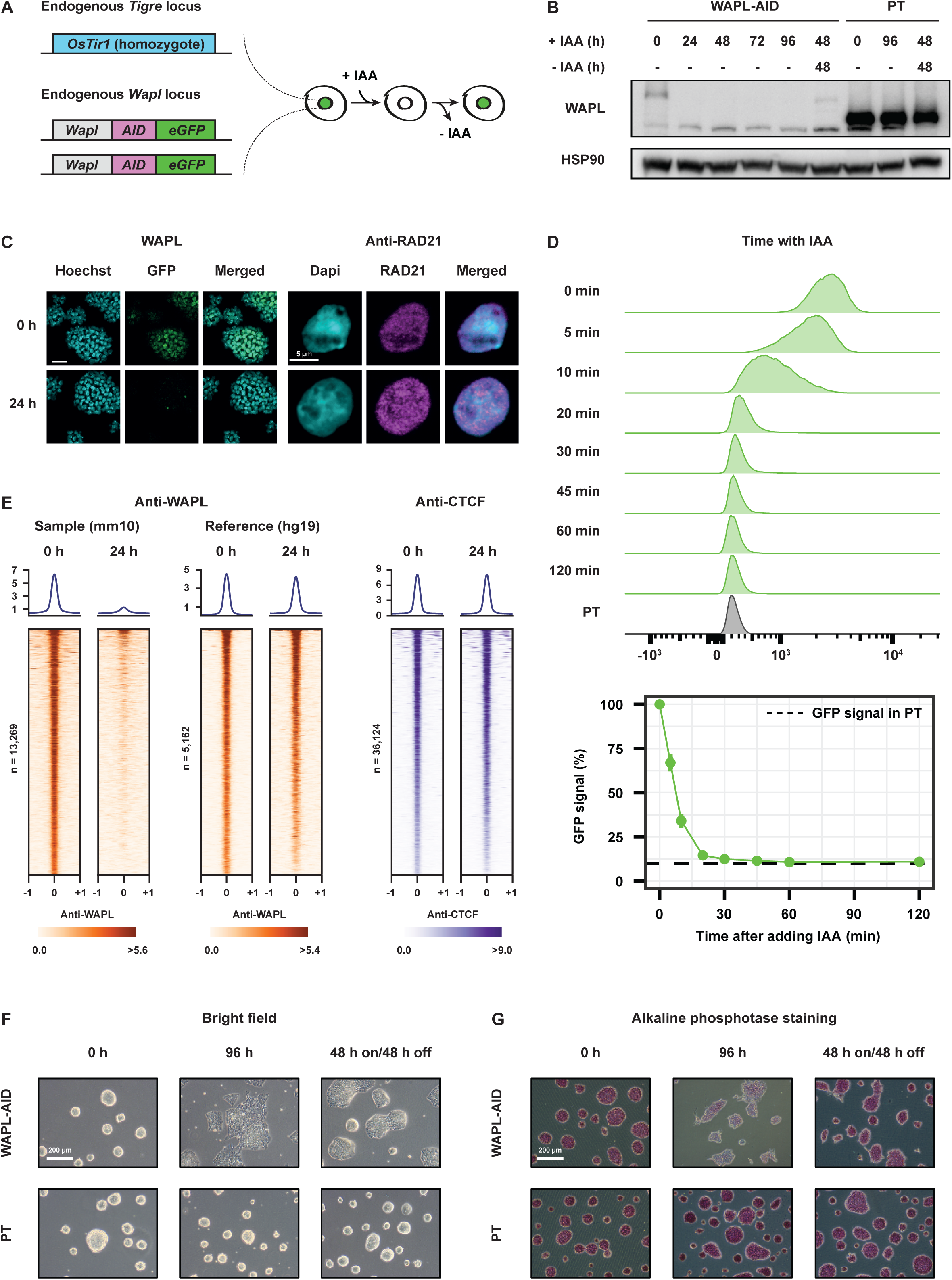
An acute degradation strategy for WAPL depletion. A) The endogenous *Wapl* gene was tagged with AID-eGFP in *OsTir1* parental mouse E14 ES cells. IAA (auxin) treatment results in rapid degradation which can be followed live through eGFP fluorescence. B) Western blot analysis of WAPL levels in WAPL-AID and parental (PT) cells following different times after IAA (auxin) treatment. C) Left panel shows live cell imaging of WAPL in untreated (0h) and treated (24h) cells. DNA is visualized with Hoechst. Right panel show immunostaining of cohesin subunit RAD21. DNA is labeled with DAPI. D) WAPL-AID-GFP levels were measured by FACS. High temporal resolution shows rapid depletion of WAPL-AID-GFP upon IAA treatment. Average signal is quantified in bottom plots. Signals are the average of three experiments. Error bars indicate standard deviation. E) ChIP-seq of WAPL and CTCF before and after IAA treatment. Heatmaps show the distribution of signal for the peaks called in the untreated cells. Average signal is shown above the heatmap. ChIP-seq data is calibrated using spiked-in human HEK-293T cells. Middle two heatmaps show WAPL ChIP-seq signal derived from the human cells. F) Bright-field microscopy images showing ES cell morphology after 96 h of IAA treatment in WAPL-AID (top row) and parental (bottom row) cells. Third column shows a wash-off experiment of 48 hours of IAA treatment followed by 48 hours of no treatment. G) Alkaline phosphatase staining is measured as a marker for pluripotency. Same treatment conditions as in F) were used.

Unexpectedly, loss of WAPL in our mESCs resulted in distinct morphological changes that are characteristic of differentiation even in 2i culture conditions (Figure 1F). The protein levels of key pluripotency transcription factors was decreased upon WAPL depletion (Figure S1C). Surprisingly, WAPL depleted cells showed a clear decrease in alkaline phosphatase staining after 4 days of IAA treatment (Figure 1G), suggesting that these cells exit the pluripotent state after WAPL degradation. We analyzed the cell cycle profiles of the control and treated cells by EdU incorporation and found no major cell cycle changes upon WAPL depletion (Figure S1D,E). Furthermore, analysis of DNA content by DAPI staining showed that no clear aneuploidy was induced in WAPL-depleted cells (Figure S1F). The morphological phenotype was fully recapitulated in a second WAPL-AID clone, also with a clear decrease of alkaline phosphatase staining intensity (Figure S1G). These data suggest that WAPL, which ensures normal off-loading of cohesin, is essential to maintain the pluripotent state of mESCs.

In order to better understand the molecular mechanisms that induce mESC differentiation following WAPL-depletion we performed RNA-seq analysis. Acute depletion of WAPL resulted in a gradual increase in transcription deregulation over the course of 96 hours IAA treatment (Figure 2A and S2A). Relatively mild effects on gene expression were observed within the first 24 hours of treatment with 330 genes showing significant changes in gene expression (FDR < 0.05, 185 up, 145 down), indicating that these genes may be directly regulated by WAPL loss (Figure 2A and S2A). We further analyzed the gene sets deregulated after WAPL depletion. Over 80% of the up-regulated biological processes (FDR < 0.01) are associated with (embryonic) tissue development, (embryonic) morphogenesis, and cell differentiation (Figure 2B). Nearly all pluripotency factors lost their normal expression after four days of WAPL depletion (Figure S2B).

**Figure 2:**
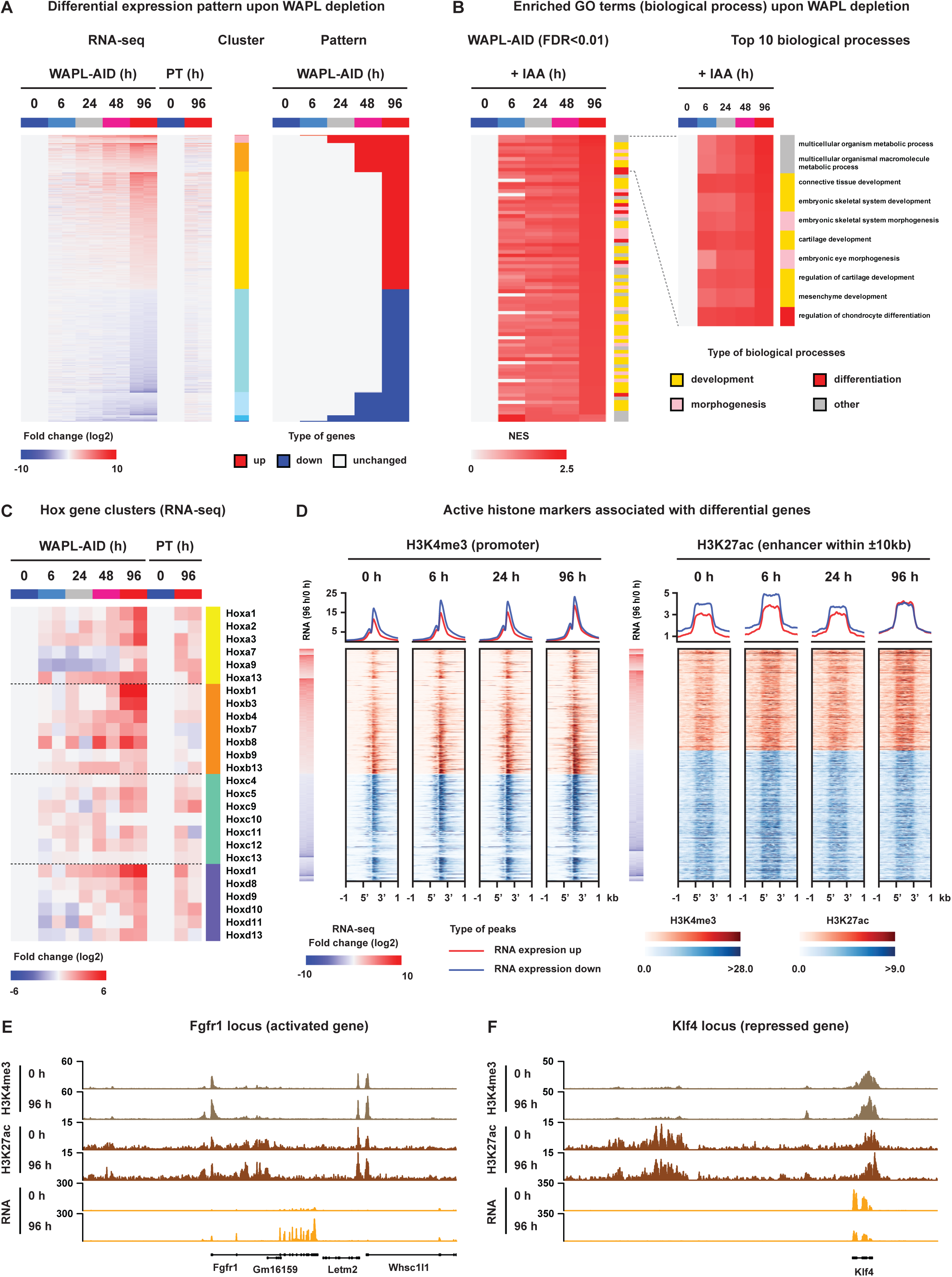
WAPL is required for maintaining a pluripotency specific transcriptional state. A) Heatmap showing the genes that are differentially expressed following IAA treatment in WAPL-AID cells. Genes are clustered according to the timepoint in which they are first observed to be differentially expressed (left panel). B) Gene Set Enrichment Analysis of RNA-seq data following WAPL depletion. Analysis was performed for Gene Ontology databases and the significant (FDR < 0.01) terms with a positive normalized enrichment score (NES) are plotted. Terms that are related to development, differentiation or morphogenesis are highlighted. C) Expression heatmap of the genes of the four *Hox* clusters. D) ChIP-seq heatmaps of two active histone modifications. Left panel shows signal of the active promoter mark H3K4me3 aligned to the promoters of differentially expressed genes. Right panel shows H3K27ac alignment to H3K27ac^+^/H3K4me3^-^ enhancers within 10kb up- or downstream of the promoter of a differentially expressed gene. E) Example region showing the H3K4me3, H3K27ac and RNA levels in the vicinity of an activated gene (*Fgfr1*) in untreated cells and cell treated for 4 days with IAA. F) Same as E), but for a repressed genes (*Klf4*).

When we further analyzed the differentially expressed genes, we found that canonical PRC2 target genes, associated with the inactive histone mark H3K27me3, showed increased expression after WAPL depletion (Figure S2C). For instance, the well-defined developmental *Hox* gene clusters (Figure 2C), which are almost uniformly covered by H3K27me3 in mESCs (Hammoud et al., 2009), were gradually up-regulated during IAA treatment. It has been suggested that loss of H3K27me3 results in increased expression of developmental genes, which in turn leads to differentiation of mESCs after WAPL knockdown (Stelloh et al., 2016). However, in mESCs cultured in 2i medium the characteristic H3K27me3 domains at canonical PCR2 target genes are absent (Joshi et al., 2015; Marks et al., 2012). We therefore profiled global H3K27me3 in our WAPL-AID cells, and the H3K27me3 occupancy showed almost no difference between the control cells and cells treated for 96 hours with IAA (Figure S2D,E). Our results suggest that activation of developmental genes is not caused by a loss of H3K27me3 mediated transcription repression.

On the other hand, gene activation is strongly associated with a gain of active promoter and enhancer histone modifications. Therefore, we profiled genome-wide H3K4me3 and H3K27ac during IAA treatment, and observed only subtle changes following 96 h of IAA treatment (Figure 2D). The active promoter mark H3K4me3 remained largely unchanged, while the active enhancer mark H3K27ac showed a very weak increase at 96 h for the activated genes (Figure 2D and 2E) but stable levels at enhancers of genes that were repressed following IAA treatment (Figure 2D and 2F). These data indicate that transcriptomic changes after WAPL depletion are not caused by massive changes in the Polycomb-repressive or enhancer-associated epigenetic landscape.

### Regions of Dynamic Cohesin are strongly enriched for pluripotency genes and enhancers

In cells lacking WAPL, cohesin rings are loaded onto chromatin, but fail to be released during interphase, leading to a global stabilization of cohesin molecules on DNA (Tedeschi et al., 2013). To determine what happens to the distribution of cohesin after acute depletion of WAPL we performed calibrated ChIP-seq (see Methods) of the core cohesin subunit RAD21. Stabilization of cohesin results in the formation of 12,554 novel cohesin binding sites. Unexpectedly, we observed a concomitant loss of 6,372 RAD21 binding sites which were clearly diminished after global stabilization of cohesin by WAPL depletion (Figure 3A). The change in cohesin binding sites suggests a global redistribution of chromatin bound cohesin upon WAPL depletion. We could recapitulate this redistribution of cohesin in an independent WAPL-AID clone (Figure S3A).

**Figure 3:**
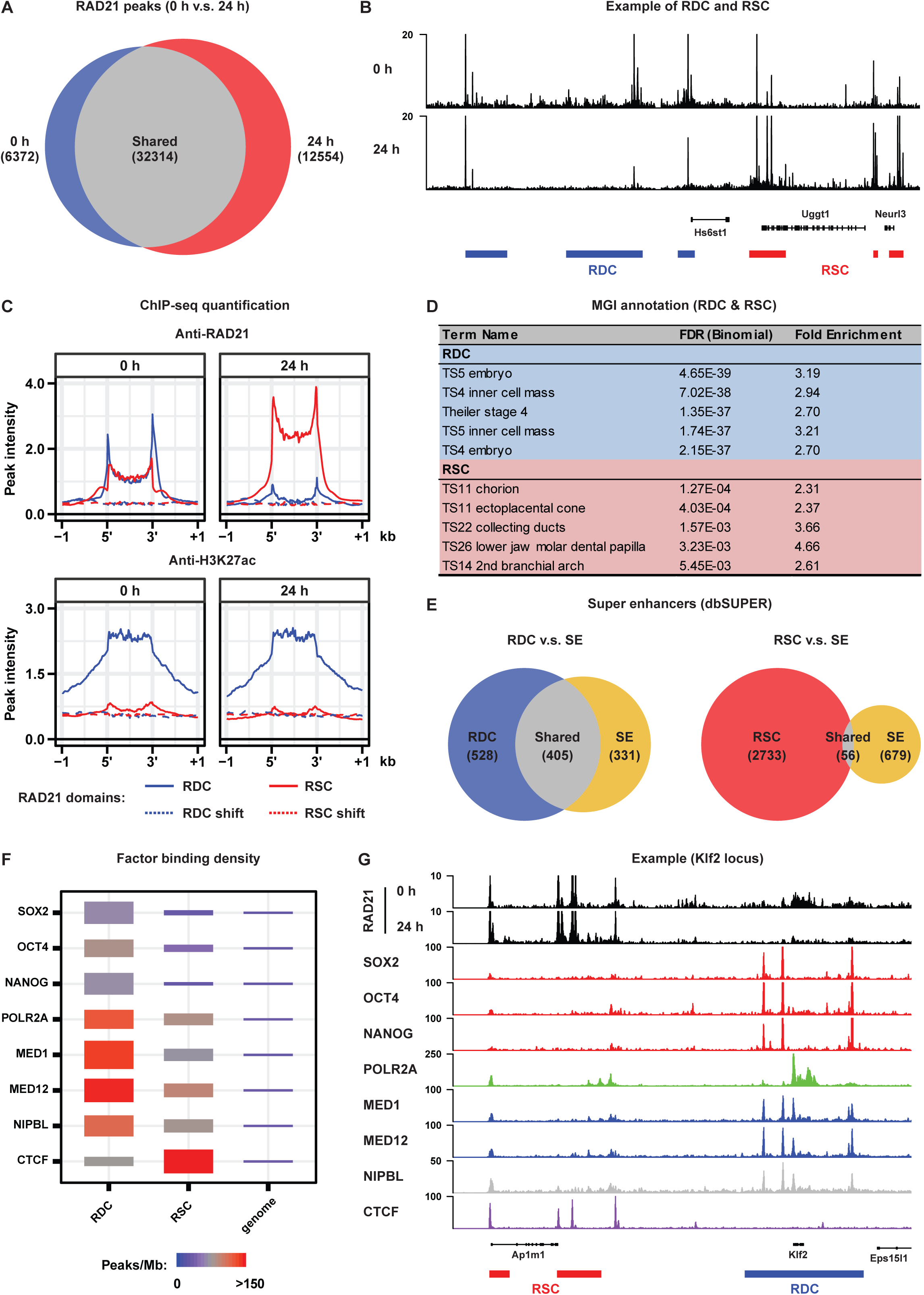
Dynamic cohesin is associated with pluripotency specific regulatory regions in mESCs. A) Venn diagram showing the shared and unique RAD21 peaks in untreated cells and cells treated for 24 hours with IAA. Peak calling was performed on a subset of 7 M sequencing reads for both of the samples. B) Example regions showing ChIP-seq coverage tracks for RAD21 in untreated and treated cells. Blue and red rectangles indicate the positions of the Regions of Dynamic Cohesin (RDC) and Regions of Stabilized Cohesin (RSC) identified by our hidden markov model. C) Top panels show alignment of RAD21 ChIP-seq data from untreated and treated cells on RDCs and RSCs. Bottom panels show alignment of H3K27ac ChIP-seq data on RDCs and RSCs. D) Table containing the top 5 (by lowest FDR) categories in a GREAT analysis(McLean et al., 2010) using the Mouse Genome Informatics (MGI) expression database for RDCs and RSCs. E) Venn diagram showing the overlap of RDCs and RSCs with super/stretch enhancers from dbSUPER (Khan and Zhang, 2016). F) DNA/chromatin binding factor density in RDCs and RSCs. Vertical width of the rectangles indicates the degree of enrichment of peak densities over the corresponding genome-wide peak densities. Color of the rectangles shows the density of the peaks per Mb. G) Example region showing the relationship between the factors analyzed in F) and the position of RDCs and RSCs.

When we looked more closely into the distribution of RAD21 in treated vs. untreated cells, we observed that RAD21 was lost over large stretches of the genome and accumulated at more focused regions (Figure 3B). To systematically analyze the lost and gained regions we developed a hidden markov model (HMM, see Methods for details), which identified 898 regions from which cohesin was lost and 2789 regions that showed increased cohesin binding after WAPL depletion (Figure S3B). Alignment of the RAD21 ChIP-seq signal on these regions clearly confirmed reduction and increase of cohesin at the lost and gained regions after WAPL depletion, respectively (Figure 3C). Next, we aligned the RAD21 ChIP-seq signal from our second independently generated WAPL-AID clone on the regions that were identified in the first clone and observed similar changes (Figure S3C). Note that these domains are not cell line or antibody specific, since alignment of publicly available ChIP-seq profiles of 5 different cohesin subunits in V6.5 mESCs at the cohesin lost and gained regions showed a similar binding pattern (Figure S3D,E). Together, these data indicate that these cohesin binding domains in mESCs are well conserved between different mouse strains. WAPL regulates cohesin turnover and is therefore essential to maintain dynamic cohesin in the nucleus. In keeping with this, we call the regions where cohesin is lost Regions of Dynamic Cohesin (RDC), to emphasize the importance of the transient cohesin binding. The loci where cohesin accumulates we will refer to as Regions of Stabilized Cohesin (RSC).

To understand the role of RDCs and RSCs we annotated the overlapping and nearby genes by performing GREAT analysis (McLean et al., 2010). In the MGI mouse developmental database (Bult et al., 2010) the RDCs show strong enrichment for genes that are expressed in early embryonic stages (Figure 3D). For the RSCs we could not find any significant gene categories that are associated with the pluripotent state (Figure 3D). When we aligned the H3K27ac ChIP-seq signal we found a strong enrichment over RDCs, but not on RSCs. (Figure 3C). This H3K27ac distribution was reminiscent of the distribution of a specific class regulatory regions called super enhancers or stretch enhancers (SE) (Parker et al., 2013; Whyte et al., 2013), which are often associated with lineage specific genes. Furthermore, it has been previously observed that SEs have a high cohesin occupancy (Cuadrado et al., 2019; Dowen et al., 2014; Ji et al., 2016). We therefore investigated how RDCs are related to SEs in mESCs. 405 out of 736 mESC SEs from dbSUPER (Khan and Zhang, 2016) overlap with RDCs (55%), for RSCs this percentage is much lower (8%, Figure 3E). In a reverse analysis we could show that the occupancy of cohesin at SEs is also depleted after WAPL depletion (Figure S3F). Moreover, we observed that the binding sites of pluripotency transcription factors SOX2, OCT4 and NANOG, active chromatin factors POL2RA, MED1 and MED12 and the cohesin loading factor NIPBL are over-represented in RDCs, while CTCF is enriched at RSCs (Figure 3F and 3G). The enrichment for components of the transcription machinery (e.g. POL2RA and Mediator subunits) at RDCs prompted us to check whether the binding of cohesin at RDCs may simply be due to the active transcriptional state. To this end, we inspected the cohesin occupancy at the promoters of highly transcribed housekeeping (i.e. non-lineage specific) genes. We found a clear enrichment of cohesin binding at the promoters of housekeeping genes that was unchanged after WAPL depletion (Figure S3G), indicating that dynamic cohesin is preferentially bound at lineage specific loci rather than to actively transcribed regions in general.

Collectively, our data show a clear global redistribution of cohesin upon WAPL depletion, leading to a loss of dynamic cohesin at lineage specific RDCs but an accumulation of stable cohesin at CTCF-dense RSCs.

### Dynamic cohesin is required to form local self-interacting domains

Cohesin is instrumental in the formation of CTCF-anchored chromatin loops and the formation of TADs (Rao et al., 2017; Schwarzer et al., 2017; Wutz et al., 2017). In order to understand the effects of cohesin redistribution on 3D genome organization, we generated Hi-C maps in control (0 h) and WAPL-depleted (24 h) cells. Contact frequency in the range of 1-10 Mb, (i.e. inter-TAD) was increased upon WAPL depletion, but decreased below 1 Mb (i.e. intra-TAD, Figure S4A). In addition, CTCF-anchored loops were extended similar to what had been observed previously (Figure S4B). These results show that acute depletion of WAPL largely recapitulates what we and others observed upon WAPL knock-out or knock-down (Haarhuis et al., 2017; Wutz et al., 2017).

When we looked at the Hi-C contact maps in regions surrounding an RDC we observed that they form regions of high self-interaction, reminiscent of TADs, for instance in the locus containing the *Sik1* gene (Figure 4A). To systematically quantify self-interaction strength on and surrounding RDCs we applied a 140 kb triangular shaped window sliding along the genome in 20 kb steps (see Figure 4B for explanation) and aligned the signal on the RDCs. We show that the high degree of self-interaction is a genome-wide feature of RDCs (Figure 4C,D), which is diminished upon WAPL depletion. In the *Klf4* locus, there are two RDCs in relatively close proximity (distance 31.9 kb, Figure 4E). Although our Hi-C data can identify general patterns we wanted to measure the effect on contact frequency at a higher resolution than can be offered by our current Hi-C data. We therefore performed high-resolution 4C-seq experiments from the promoter of the *Klf4* gene (Figure 4F). WAPL depletion leads to a clear decrease in contact frequency between the two RDCs. Similar results were obtained for the *Klf9* locus (Figure S4C,D). RSCs on the other hand show a lower degree of self-interaction (Figure 4C,D). We also wanted to check how RDCs and RSCs are related to TAD boundaries. To this end we calculated the insulation score, which is a measure for how strongly two genomic regions are segregated in a Hi-C contact map. We found that RSCs, in contrast to RDCs, have a low insulation score, indicating strong insulation between neighboring genomic regions (Figure S4E). This is consistent with the enrichment of CTCF in RSCs, which acts as a boundary protein (Nora et al., 2017). In summary, these data show that the redistribution of cohesin affects the local chromatin interactions without changing the position of TAD boundaries across the genome.

**Figure 4:**
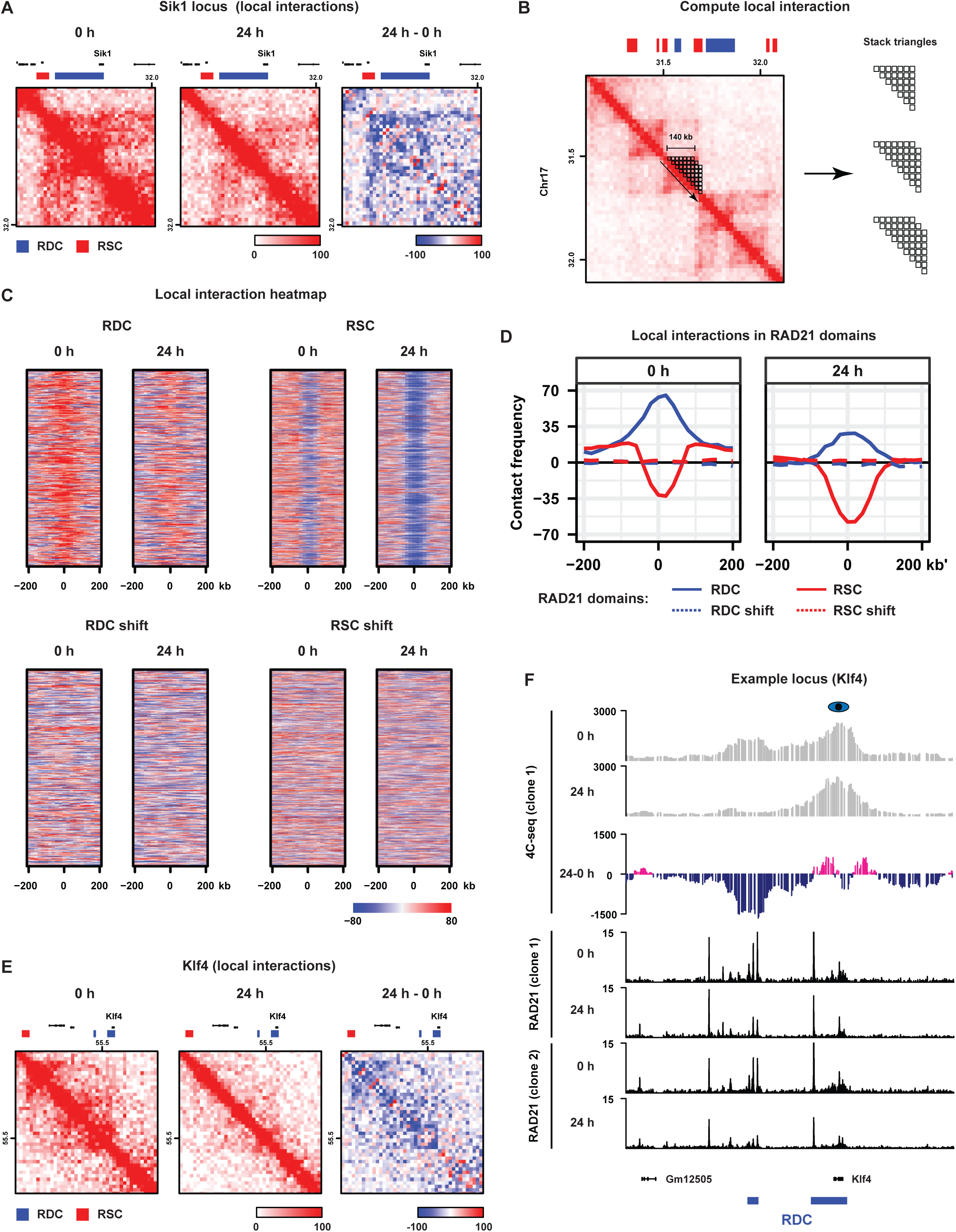
Dynamic cohesin creates regions of increased self-interaction. A) Hi-C data and RDC and RSC locations shown for the *Sik1* locus. Third panel shows differential Hi-C contacts between treated and 24h IAA treated cells. B) The self-interaction (SI) score is calculated by averaging the contact frequency within a triangle off the diagonal. The triangle is moved along the chromosome in steps of one Hi-C bin. The resulting SI score is aligned to the RDCs and the RSCs. C) Heatmaps show the SI scores for RDCs and RSCs and for randomly shifted RDC and RSC position. D) Average SI scores for RDCs and RSCs in untreated cells (left panel) and in cells treated for 24 hours with IAA (right panel). E) Same as A) but for *Klf4*. F) High-resolution 4C-seq data (see Methods) for the *Klf4* locus. Viewpoint primers were designed as close as possible to the *Klf4* promoter. Top two rows show the contact profile for the *Klf4* promoter (average of two template preparations from two depletion experiments). Third row shows the differential contact frequency between 0h 4C profile and the 24h IAA treated 4C profile. Bottom rows show ChIP-seq data for RAD21 in the *Klf4* locus from two WAPL-AID clones for treated and untreated cells.

### Cohesin is required for maintaining pluripotency-specific gene expression

Next, we wanted to know how the redistribution of cohesin affected gene expression. To this end we determined the enrichment of differentially regulated genes on RDCs and RSCs. We found that down-regulated genes are specifically enriched among the genes that are nearest to an RDC (Figure 5A) and include genes such as *Tfapc2* and *Tet2*. This enrichment was observed across all the time points after WAPL depletion, suggesting that the effect is a direct result of a loss of cohesin in these regions. For the RSCs there is no significant enrichment of down-regulated genes (Figure S5A) and for up-regulated genes we found enrichment for neither RDCs or RSCs (Figure 5A, S5A). To test whether the downregulation is a result of diminished pluripotency transcription factor binding or Mediator binding, we performed ChIP-seq for NANOG, SOX2, and OCT4, as well as a core subunit of the Mediator complex MED1. WAPL depletion did not affect the binding of any of these proteins (Figure 5B, S5B), suggesting an alternative regulatory mechanism. In order to test the effect of WAPL depletion on promoter-enhancer contact frequency we performed high-resolution 4C analysis. We designed a viewpoint on a NANOG binding site in an RDC downstream of the *Sik1* locus. We found that there was a ∼9-fold decrease in contact frequency after 24 hours of WAPL depletion (Figure 5C), and an expression loss of the *Sik1* gene. A similar effect was observed at the *Elf3* locus, with a clear decrease of local chromatin interactions and expression of the *Elf3* gene after WAPL depletion (Figure S5C). These results suggest that the binding of dynamic cohesin is crucial for the maintenance of expression through the maintenance of promoter-enhancer interactions.

**Figure 5:**
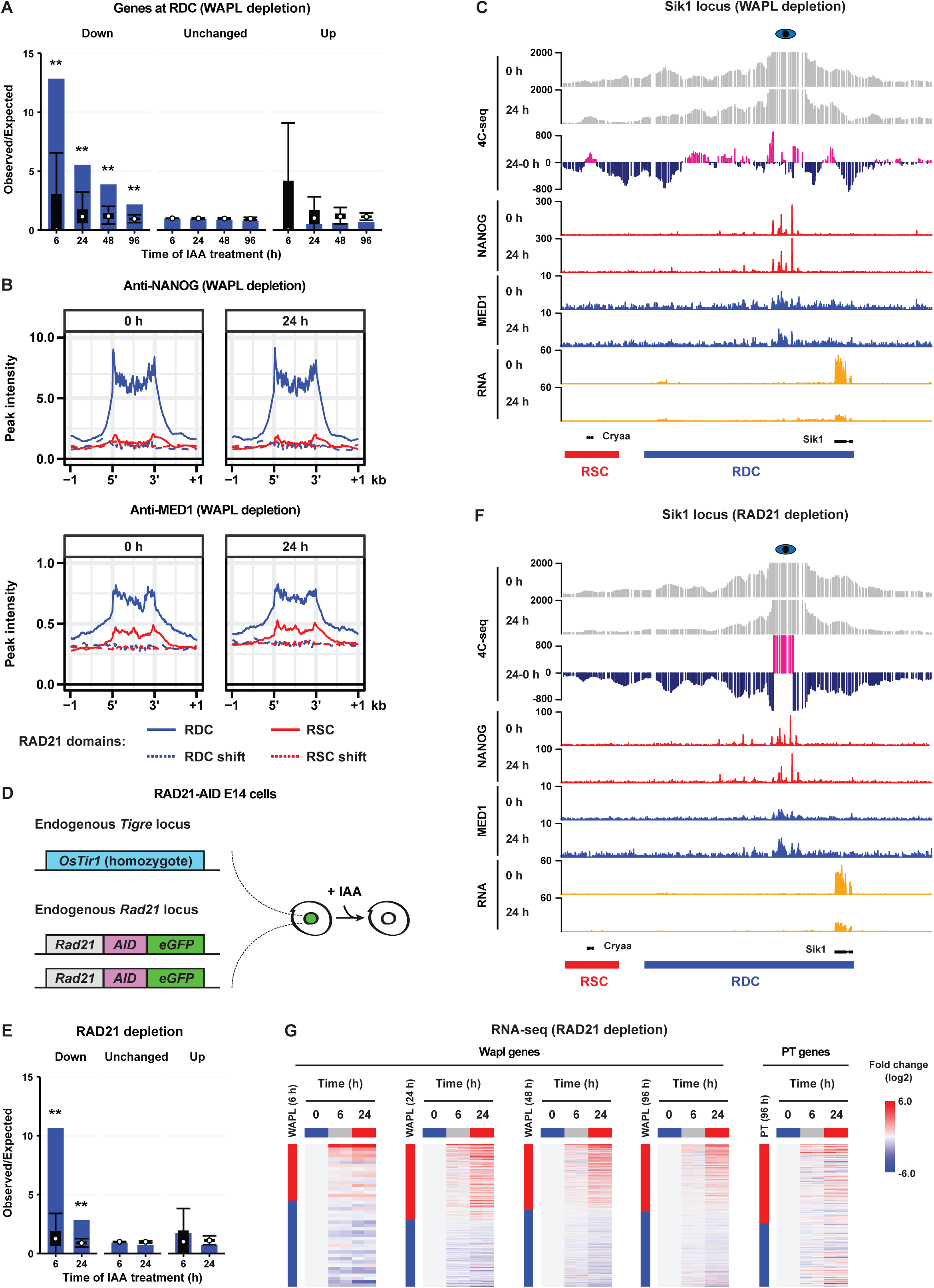
Dynamic cohesin is required for maintaining pluripotency specific expression. A) For every RDC the closest gene was identified and the observed over expected ratios (see Methods) for genes that were downregulated, upregulated or unchanged were determined. Black boxes show interquartile range, whiskers the 5th and 95th percentiles and white dot the median of 10.000 circular permutations. B) Average profiles of NANOG (top) and MED1 (bottom) binding over RDCs and RSCs characterized by ChIP-seq before and after WAPL depletion. C) High-resolution 4C-seq analysis of the *Sik1* locus. Viewpoint primers were designed close to a *Sik1* distal NANOG binding site. 4C data is visualized as in Figure 4F. D) The endogenous *Rad21* gene was tagged with AID-eGFP in *OsTir1* parental mouse E14 ES cells. E) Enrichments scores of unchanged, up- or down-regulated genes in RAD21 depleted cells after 6 or 24 hours of IAA treatment calculated and plotted similar to A). F) 4C and ChIP-seq for the *Sik1* locus similar as in C) but for the RAD21 depletion line. G) For the differentially expressed genes detected in WAPL-AID cells, the relative expression levels in RAD21 depleted cells are plotted as heatmaps. The vertical bars show the differentially expressed genes in the respective timepoints following WAPL depletion. Red signifies upregulated genes, blue down. Next to the bar the RAD21 depletion time series heatmap for the same genes is shown.

We reasoned that if local depletion of cohesin from the RDCs results in decreased expression of the RDC genes, we should be able to phenocopy this by a complete loss of cohesin. To test this, we generated a degron line to acutely deplete RAD21. We fused AID-GFP in frame with RAD21 (Figure 5D). The AID-tagged RAD21 protein was completely degraded after 6 hours of IAA treatment (Figure S5D). After 24 hours of IAA treatment, RAD21-AID cells showed a similar morphological change as WAPL depleted cells (Figure S5E). In addition, Western blot analysis showed a clear diminishment of the key pluripotency factors OCT4 and NANOG (Figure S5F). After 6 and 24 hours of RAD21 depletion RNA-seq analysis revealed 218 (82 up and 136 down) and 4,144 (2,176 up and 1,968 down) differential genes, respectively. We intersected the RNA-seq data that we generated in RAD21-AID cells with RDC associated genes and found that there was an enrichment for down-regulated genes, but not upregulated genes (Figure 5E). Like in the WAPL depleted cells, the *Sik1* gene also showed a decrease in expression upon RAD21 depletion. As expected, 4C analysis in the *Sik1* locus showed that contact frequency between the *Sik1* promoter and its distal regulatory elements was decreased after 24hrs of RAD21 depletion (Figure 5F). Again, this cannot be explained by a loss of NANOG or MED1 binding, because ChIP-seq of these factors revealed little difference upon RAD21 depletion (Figure 5F, S5I). Importantly, expression changes upon RAD21 depletion were strikingly similar to expression changes as a result of WAPL depletion (Figure 5G). A similar effect on chromatin interaction and gene expression was seen for the *Elf3* locus in the RAD21 depletion experiment (Figure S5J).

In summary, our data suggest that dynamic cohesin binding at RDCs is essential to control expression of a subset of genes in mESCs. Loss of cohesin binding in these regions, either via redistribution of cohesin as a result of stabilization of the complex or the complete loss of cohesin leads to decreased expression of genes associated with RDCs without altering pluripotency specific transcription factor binding.

### OCT4 creates a platform for cohesin binding

CTCF-independent cohesin binding at lineage specific sites has been reported for a number of cell types (Faure et al., 2012; Kagey et al., 2010). However, how cohesin is recruited to these sites remains unclear. Based on our observation that dynamic cohesin is found at the binding sites of pluripotency transcription factors we hypothesized that some of these factors are responsible for the binding of cohesin molecules at these cell-type specific regulatory regions. To test this hypothesis, we employed a published OCT4-FKBP cell line (Boija et al., 2018) and generated a NANOG-FKBP cell line (Figure 6A). FKBP fusion proteins can be rapidly degraded by addition of the heterobifunctional dTAG molecule (Nabet et al., 2018). Nearly complete depletion of OCT4 and NANOG was achieved within 24 h of adding 500 nM dTAG-13 molecule into the cell culture (Figure 6B).

**Figure 6:**
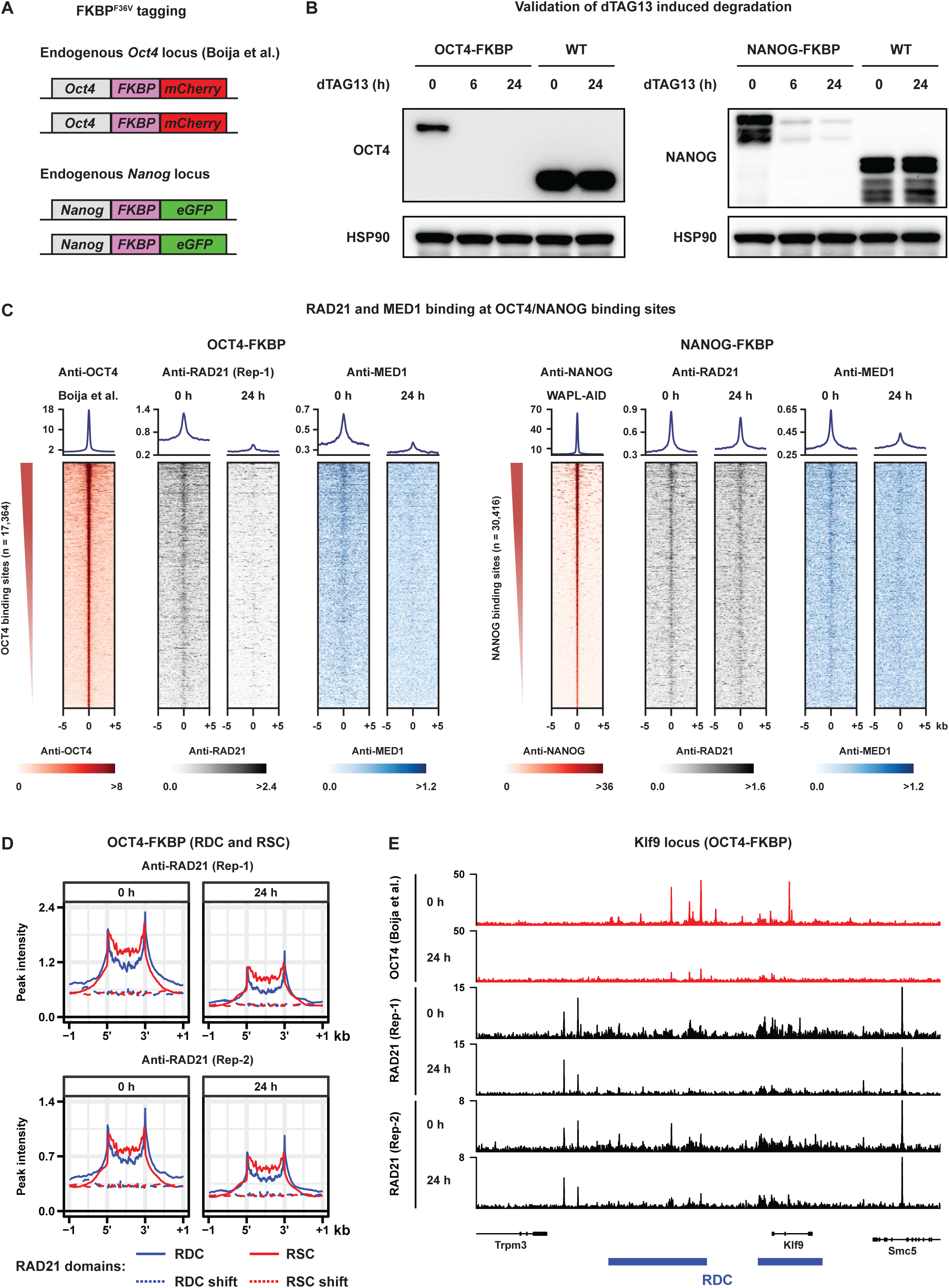
A subset of cohesin binding is dependent on the transcription factor OCT4. A) Two FKBP tagged cell lines were used, a previously published OCT4-FKBP-mCherry line and a NANOG-FKPB-eGFP (see Methods for details on construction). The FKBP degron can be degraded with the dTAG-13 molecule (Nabet et al., 2018). Western blot shows protein levels of OCT4 (left) and NANOG (right) after 6 and 24 hours of dTAG-13 treatment. C) ChIP-seq heatmaps showing RAD21 and MED1 levels over OCT4 binding sites (Boija et al., 2018) following 24h of OCT4 depletion (left panel); ChIP-seq heatmaps showing RAD21 and MED1 levels over NANOG binding sites identified in the WAPL-AID cell line following 24h of NANOG depletion (right panel). D) RAD21 ChIP-seq alignment over RDCs and RSCs following OCT4 depletion for two independent replicates. E) Example locus containing the *Klf9* gene shows the binding of RAD21 over RDCs in treated and untreated OCT4-FKBP cells. OCT4 ChIP-seq track shows previously published OCT4 profiles in the same cell line (Boija et al., 2018).

Next, we examined what happened to RAD21 binding at OCT4 and NANOG binding sites before and after dTAG-13 treatment. Strikingly, OCT4 depletion resulted in a strong decrease of cohesin binding at OCT4 binding sites, while NANOG depletion did not affect cohesin binding at NANOG binding sites (Figure 6C). However, cohesin binding at CTCF sites is largely unchanged following OCT4 or NANOG depletion (Figure S6B). As expected, cohesin occupancy at the RDCs was decreased in the OCT4 but not in the NANOG depletion experiment (Figure 6D,E, S6D). It has been shown that the pluripotency transcription factors OCT4 and ESRRB are responsible for recruiting the Mediator complex to chromatin (Boija et al., 2018; Sun et al., 2019). To confirm that we have functional depletion of both OCT4 and NANOG in their respective degron lines we examined MED1 binding before and after depleting OCT4 and NANOG. We found that both OCT4 and NANOG depletion led to a decrease in MED1 binding at OCT4 and NANOG binding sites, respectively (Figure 6C). Collectively, these data show that cohesin binding to pluripotency-specific regulatory sites is dependent on OCT4 but not NANOG.

### Dynamic cohesin binding sites are found in differentiated cells

Following the identification of RDCs in mESCs we wondered whether the redistribution of cohesin was unique to pluripotent cells or could be found in other cell types as well. To address this question, we differentiated WAPL-AID mESCs into neural progenitor cells (NPCs) *in vitro* following a standard differentiation protocol (Figure 7A) (Peric-Hupkes et al., 2010). We confirmed that the generated NPCs were positive for NESTIN (an NPC marker) and negative for GFAP (an astrocyte marker) (Figure 7B, S7A), and did not show alkaline phosphatase staining (Figure S7B). The differentiated cell line still expressed the WAPL-AID-GFP fusion protein and treatment with IAA effectively degraded the protein (Figure 7C). In mESCs, WAPL depletion leads to a loss expression of lineage-defining (i.e. pluripotency) genes, in NPCs IAA treatment led to loss of the NPC marker NESTIN in the WAPL-AID NPCs but not parental cells (Figure 7B, S7A), indicating that WAPL depletion results in loss of neural progenitor identity.

**Figure 7:**
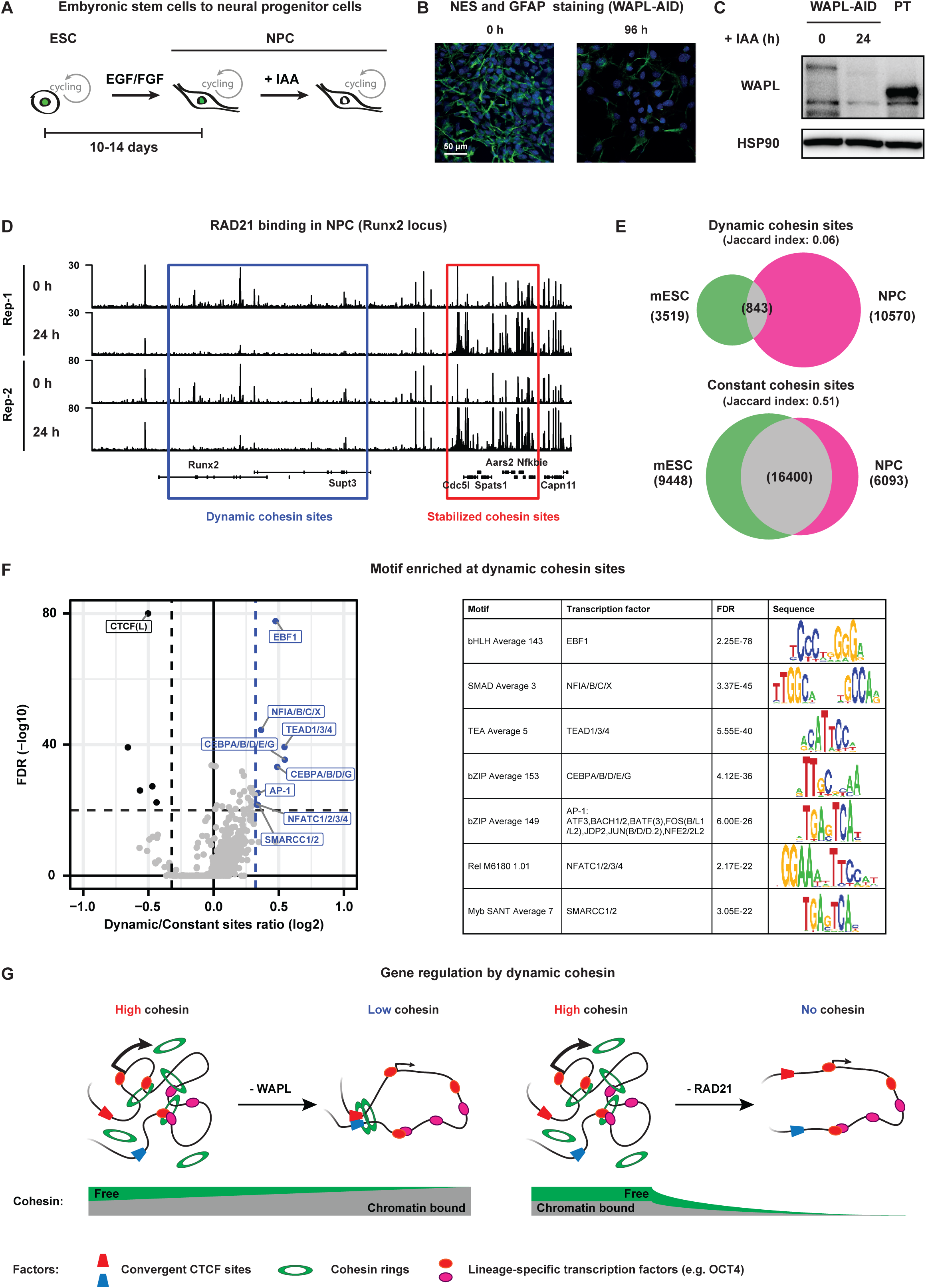
Dynamic cohesin binding sites are a feature differentiated cells. A) WAPL-AID cells were differentiated to neural precursor cells using a standard differentiation protocol (Peric-Hupkes et al., 2010). Immunofluorescence of neuronal markers NESTIN and GFAP (green: NESTIN, red: GFAP, blue: DAPI) for treated and untreated neural precursor cells. C) Western blot analysis of WAPL in untreated cells and 24 hours IAA treated cells. Parental (PT) NPCs, derived from *OsTir1* parental ES cells were taken along as control. D) Example region shows the ChIP-seq tracks of RAD21 for treated and untreated WAPL-AID NPCs. E) Venn diagram showing overlap of dynamic and constant cohesin binding sites of mESCs and NPCs. Dynamic cohesin sites are regions bound in the presence of WAPL, but not bound in the absence of WAPL. Constant cohesin sites are regions bound in the presence and absence of WAPL. Peak calling was performed on a subset of 7 M sequencing reads for all the samples. F) Motif enrichment analysis for dynamic cohesin binding sites. Sites bound preferentially in untreated cells (“Dynamic”) and sites bound stably in both treated and untreated cells (“Constant”) were analyzed with GimmeMotifs (van Heeringen and Veenstra, 2011). P-values are calculated using the Fisher exact test for motif frequency for dynamic and constant peaks for every motif. Log_2_ fold-change of motif frequency is determined by calculating the ratio of relative motif frequency (i.e. corrected for the total motif frequency of all motifs) between dynamic and constant peaks (see Methods for details on fold-change and p-value calculation). G) Model for the consequences of WAPL and RAD21 depletion. WAPL depletion results in an accumulation of cohesin at CTCF binding sites. RAD21 depletion leads to a general loss of cohesin from the genome. Both WAPL and RAD21 depletion results in loss of cohesin binding dynamics from lineage-specific transcription factor binding sites and decreased 3D genome interactions between promoters and these sites.

To determine whether cohesin is redistributed after WAPL depletion in NPCs, we performed RAD21 ChIP-seq in NPCs with and without IAA treatment. We found that cohesin stabilization resulted in a clear redistribution (Figure 7D). In control and WAPL depleted NPCs we found a total of 11413 and 10591 RAD21 binding sites unique to either condition, respectively, across two replicates (Figure S7C). We found 22644 RAD21 binding sites that were found in treated and untreated cells (i.e. ‘constant’ binding sites). When we compared the constant RAD21 binding sites that were identified in both mESCs and NPCs we found a strong overlap in the binding sites (Jaccard index 0.51, Figure 7E). However, when we performed the same analysis for sites that were lost upon WAPL depletion in mESCs and NPCs a much weaker overlap was observed (Jaccard index 0.06, Figure 7E). In order to annotate these dynamic cohesin sites, we performed a stringent identification using DESeq2 (see Methods, Figure S7D). We subsequently performed motif analysis (see Methods) to identify potential transcription factors associated with dynamic cohesin binding sites. As expected, the constant sites show a strongly significant enrichment for the CTCF motif (Figure 7F). For the cohesin binding sites lost after WAPL depletion we observed a significant enrichment of the transcription factors that can be associated to neuronal development, such as EBF1 (Garel et al., 1999) and nuclear factor I (NFI) (Driller et al., 2007). These results show that stable CTCF associated cohesin sites are largely tissue-invariant, but that dynamic cohesin sites are lineage specific, associated with lineage specific transcription factors and are likely to be involved in the control of cellular identity.

## DISCUSSION

### Dynamic cohesin is crucial in maintaining lineage specific expression

In this study, we used actute depetion of chromatin-associated proteins to study the role of the cohesin complex in the regulation of lineage specific genes. Importantly, the effects on gene expression following stabilization of cohesin by WAPL depletion can be almost phenocopied by acute depletion of RAD21. These paradoxical results can be explained by considering that it is the dynamic fraction of the nuclear cohesin pool that is important for the regulation of genes. Indeed, the regions where cohesin is lost after cohesin stabilization and should *ipso facto* be binding sites for dynamic cohesin are nearest to down-regulated genes. This is corroborated by the observation that the RDCs are enriched for active enhancer marks and pluripotency transcription factors. Our 3D genome analyses suggest that dynamic cohesin mediates interactions between promoters and enhancers and that disruption of these contacts by either WAPL or RAD21 depletion leads to a decrease in expression. This activating role is consistent with the observation that in mature mouse macrophages inducible knock-out of *Rad21* resulted in a failure to upregulate genes upon stimulation with LPS (Cuartero et al., 2018). Moreover, although in human HCT116 colon cancer cells there was a relatively mild effect on gene expression following acute RAD21 depletion, there was an enrichment of downregulated genes closer to SEs. Given that the RDCs that we identified cover the majority of SEs, these results are consistent with our data. Consistent with our observation that loss or redistribution of cohesin results in differentiation, heterokaryon mediated reprogramming fails when in the absence of RAD21 (Lavagnolli et al., 2015).

We would like to note that acute depletion of RAD21 in the study of gene regulation is challenging due to the essential role cohesin also plays in sister chromatid cohesion (Nasmyth and Haering, 2009) and DNA repair (Strom et al., 2007). With the importance of dynamic cohesin in the regulation of expression, WAPL actually serves as an ideal proxy for modulating cohesin’s role in gene expression. The fact that WAPL depletion also shows a loss of non-CTCF cohesin sites in differentiated cells shows that this regulatory axis can be exploited for studying the role of cohesin in gene regulation beyond mESCs.

### A role for loop extrusion in gene regulation?

We and others have previously shown that stabilization of cohesin results in increased loop lengths (Haarhuis et al., 2017; Wutz et al., 2017). We fully recapitulate this phenotype using acute depletion of WAPL in mESC. We also observe that regions where cohesin accumulates are enriched for the boundary protein CTCF. The loop extension phenotype after WAPL depletion is in line with the loop extrusion model that has been proposed to explain TAD formation and the convergent orientation of CTCF sites forming loops (Fudenberg et al., 2016; Sanborn et al., 2015). The extrusion model posits that formation of TADs is dependent on a cycle of loading, extrusion and off-loading (Fudenberg et al., 2016). Stabilization of cohesin breaks this cycle and results in diminished intra-TAD interactions (Haarhuis et al., 2017). Here we have identified regions of dynamic cohesin, that form self-interaction domains. Depletion of WAPL results in decreased self-interaction in these domains and a decreased contact frequency between promoters and regulatory elements.

In keeping with the above, we believe that the extrusion cycle, which depends on dynamic cohesin, is important for bringing distal regulatory sites into contact with their cognate promoter as well. Loss of dynamic cohesin by either removing all cohesin molecules (RAD21 depletion) or exhausting the freely available cohesin (WAPL depletion), disrupts the loop extrusion cycle. It is important to emphasize that compared to a diffusion model, loop extrusion effectively turns a 3D search into 1D scanning (Bulger and Groudine, 2011). Furthermore, the diffusion model hypothesizes that promoter-enhancer interactions within a TAD are mediated by high local concentration of diffusible activators (Gurumurthy et al., 2019), such as transcription factors and the Mediator complex. Although, the transcription factor and Mediator binding is largely unchanged in the context of WAPL and RAD21 depletion, the contact frequency between promoters and enhancers is strongly diminished, emphasizing the importance of cohesin over diffusion mediated interactions. Although, the details of cohesin mediated loop extrusion remain to be worked out, it is clear that we have identified a subset of genes that depends on cohesin-mediated promoter-enhancer communication for their activation.

### Cohesin binding to pioneer factor binding sites

Cohesin has previously been shown to overlap with the binding sites of sequence specific transcription factors (Faure et al., 2012; Nitzsche et al., 2011). In mESCs we found a subset of weakly bound cohesin sites overlapping with binding sites of pluripotency factors OCT4, SOX2 and NANOG. Stablization of cohesin results in loss of cohesin from these binding sites. This binding can be either the result of direct or indirect recruitment by transcription factors at these sites or the result of stalling of the extrusion process akin to CTCF. Co-immunoprecipitation (Co-IP) experiments have identified an interaction between OCT4 and SMC1A in mESC (van den Berg et al., 2010), which could indicate that cohesin is directly recruited by OCT4, although the interaction between OCT4 and cohesin may occur via a third protein that interacts with both. Note, that Co-IP experiments for NANOG picked up an interaction with cohesin subunit STAG1 (Nitzsche et al., 2011), whereas, NANOG depletion did not affect cohesin binding in our experiments. Alternatively, OCT4, which acts as a pioneer factor, creates regions of open chromatin in conjunction with the chromatin remodeler BRG1 (King and Klose, 2017). Since cohesin is recruited to sites of open chromatin (Lopez-Serra et al., 2014), this could explain why loss of OCT4, but not NANOG, leads to a loss of cohesin binding. However, not all open chromatin sites are enriched for cohesin binding, suggesting that there are likely additional signals to bring cohesin specifically to OCT4 bound open chromatin sites. Although the stalling scenario is a formal possibility to explain why cohesin is bound to transcription factor binding sites, this would effectively mean that transcription factor binding sites act as boundaries, for which there is currently little evidence in mammalian cells.

Importantly, we found that there is no difference in NANOG binding upon either WAPL or RAD21 depletions. This means that cohesin is not required for maintaining an open chromatin structure to allow for transcription factor binding in mESCs (Yan et al., 2013). Our results also seem to conflict with earlier observations, where heterozygous knock-out of *Rad21* leads to a loss of transcription factor binding (Faure et al., 2012). However, the number of lost sites is rather limited and the differences may be attributed to pleiotropic effects of pan-cellular knock-out of one of the *Rad21* alleles. In our acute depletion experiments we can assay the direct effects on transcription factor binding and we do not find a severe change in binding. From this we conclude that cohesin is not involved in transcription factor recruitment.

Our results shed light on the mechanism by which cohesin is involved in regulating gene expression. We show that a subset of cohesin binding sites depend on the activity of WAPL and are largely independent of CTCF and CTCF-anchored loops. Rather the binding of cohesin depends on a specific subset of transcription factors. The pioneer transcription factor OCT4 can create an open chromatin region which may serve as a binding platform for cohesin. Through the loop forming capacity of the cohesin complex regulatory elements may in this way be connected to promoters in a dynamic manner to enhance expression. Dynamic cohesin binding sites are often found in proximity of lineage specific genes emphasizing the importance of this complex for the proper expression of genes throughout development and may explain the pleiotropic effects found in cohesinopathies such as Cornelia de Lange syndrome (Krantz et al., 2004).

## ACKNOWLEDGEMENTS

We thank the NKI Genomics Core Facility for help with sequencing, the NKI Bioimaging Facility for help with microscopy, the NKI Flow Cytometry Facility for help with single cell sorting of genome edited cells. We thank Masato Kanemaki for his suggestions on AID tagging and sharing *OsTir1* antibody. We thank Behnam Nabet and Nathaniel Gray for providing the dTAG-13 molecule. We thank Richard Young for providing the OCT4-dTAG ES cell line. Work in the de Wit lab is supported by an ERC StG 637587 (‘HAP-PHEN’) and a Vidi grant from the Netherlands Scientific Organization (NWO, ‘016.16.316’). N.Q.L. is supported by a Veni grant from the Netherlands Scientific Organization (NWO, ‘016.Veni.181.014’). N.Q.L., M.M., T.v.d.B, L.B., H.T., M.S., and E.d.W. are part of Oncode which is partly financed by the Dutch Cancer Society.

## AUTHOR CONTRIBUTIONS

N.Q.L. and E.d.W. conceived and designed the study; N.Q.L., M.M., L.B., and H.T. performed experiments in the lab of E.d.W.; E.P.N. engineered *OsTir1* and RAD21-AID cell lines in the lab of B.G.B; N.Q.L., T.v.d.B., M.S. and E.d.W. analyzed data; E.d.W. supervised the study; N.Q.L. and E.d.W. wrote the manuscript with input from all authors.

## STAR★Methods

### Key Resources Table

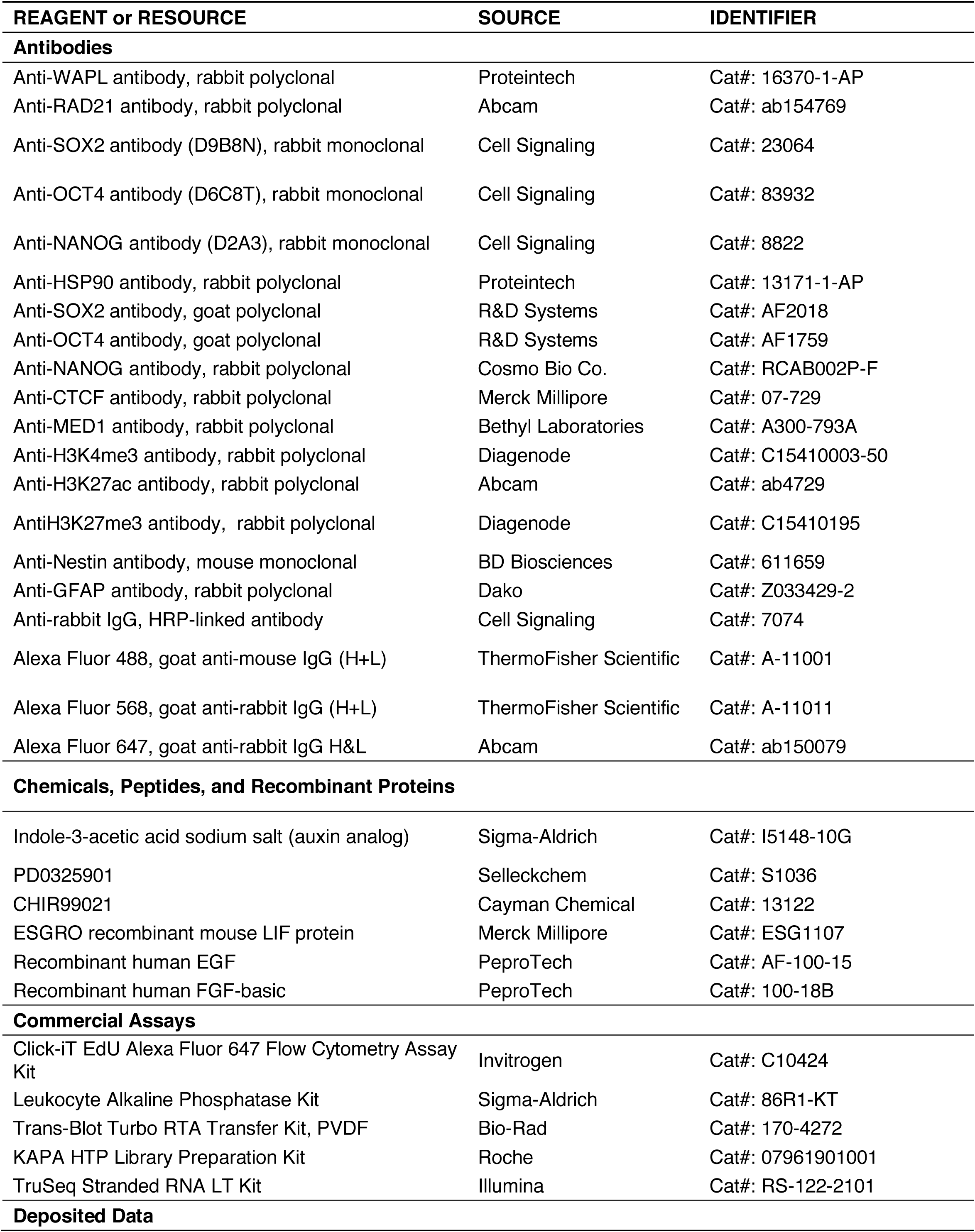
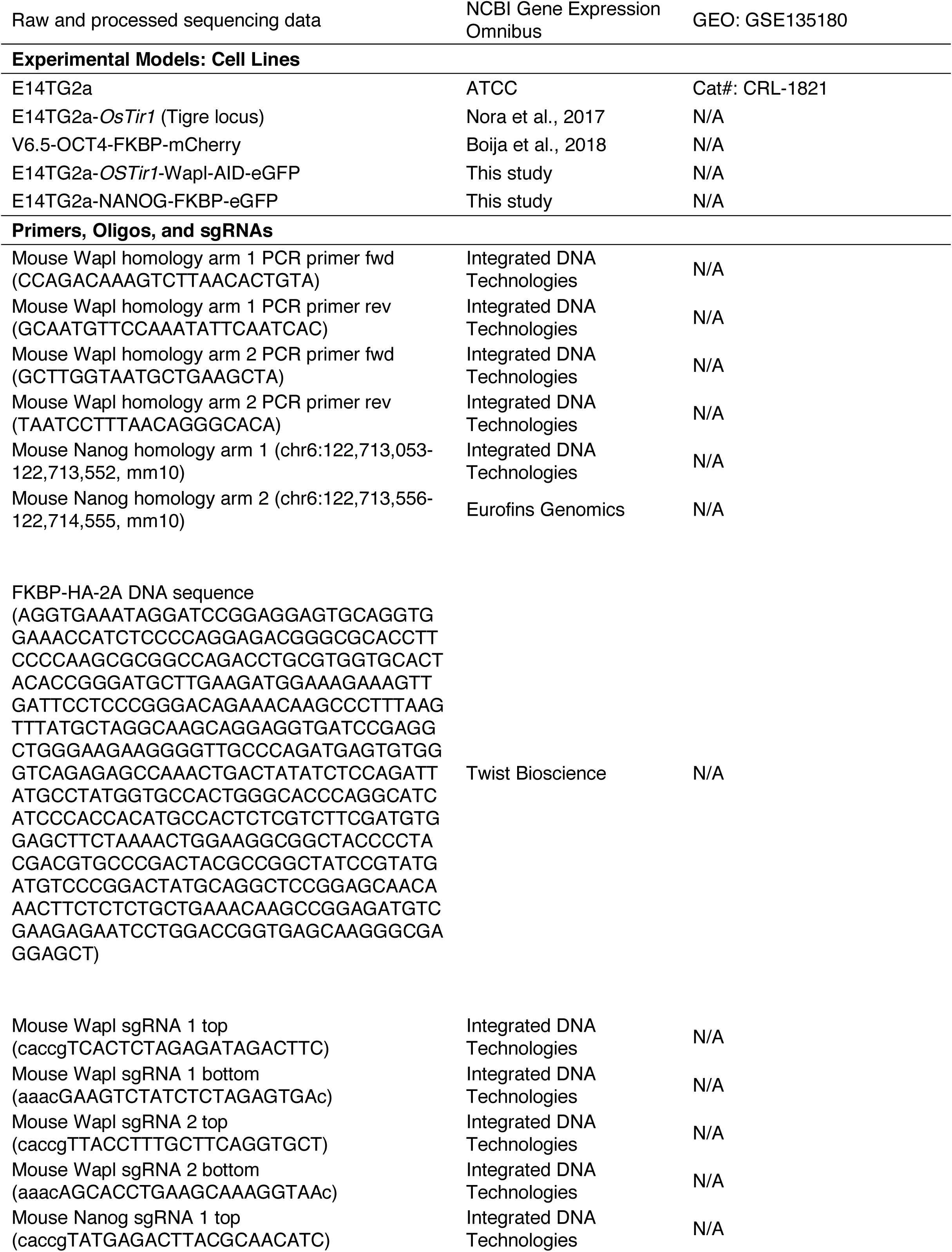
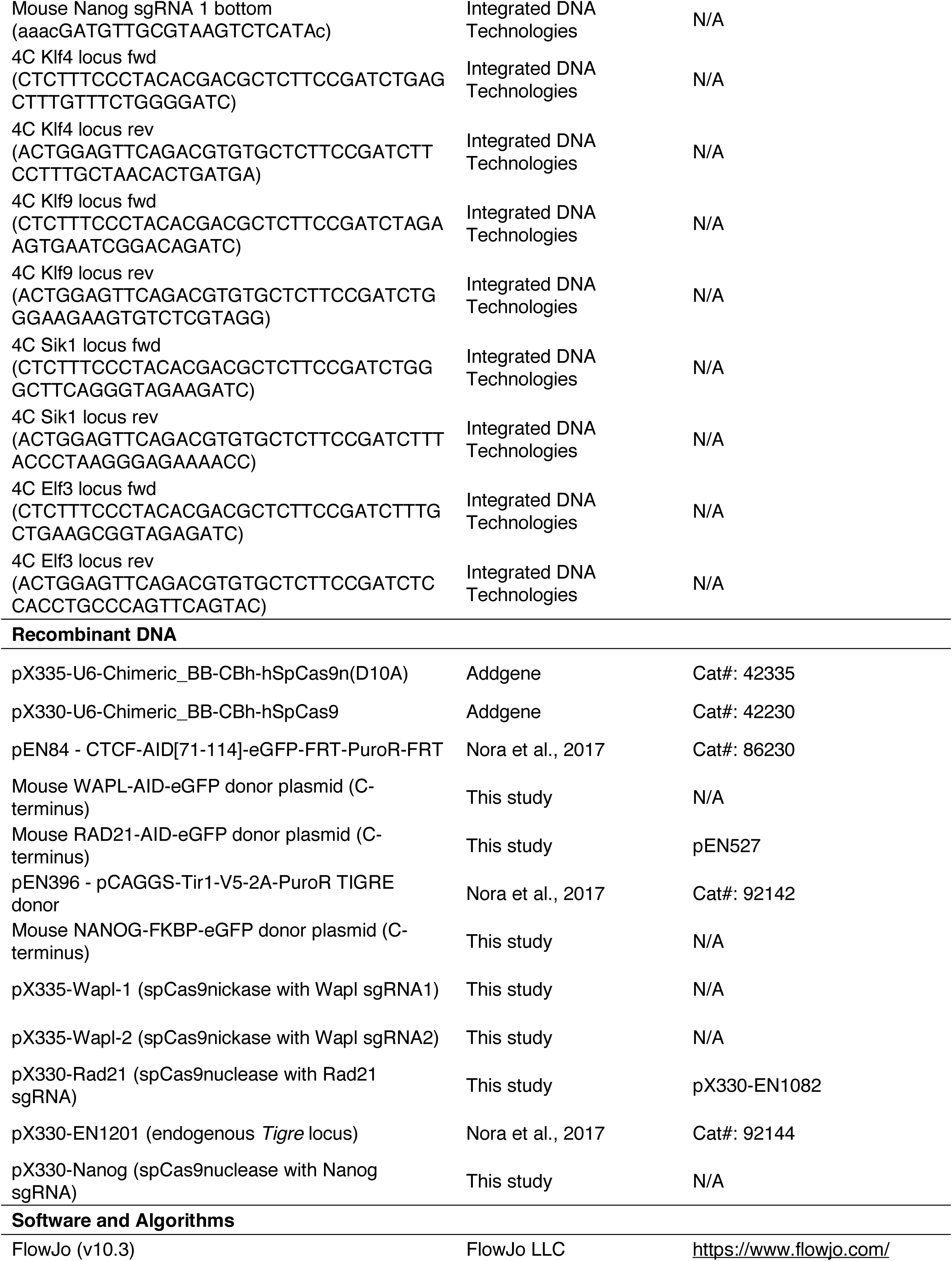
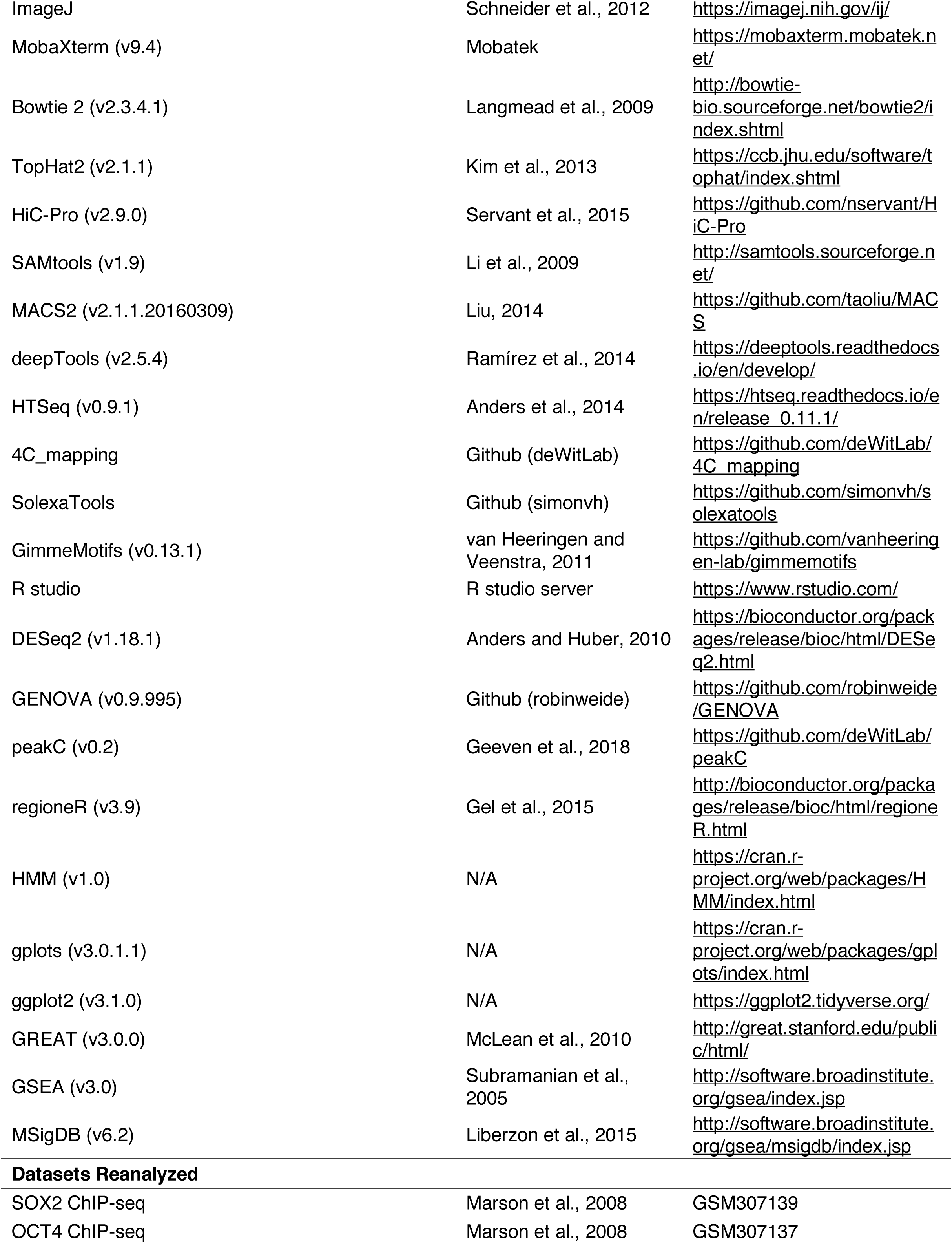
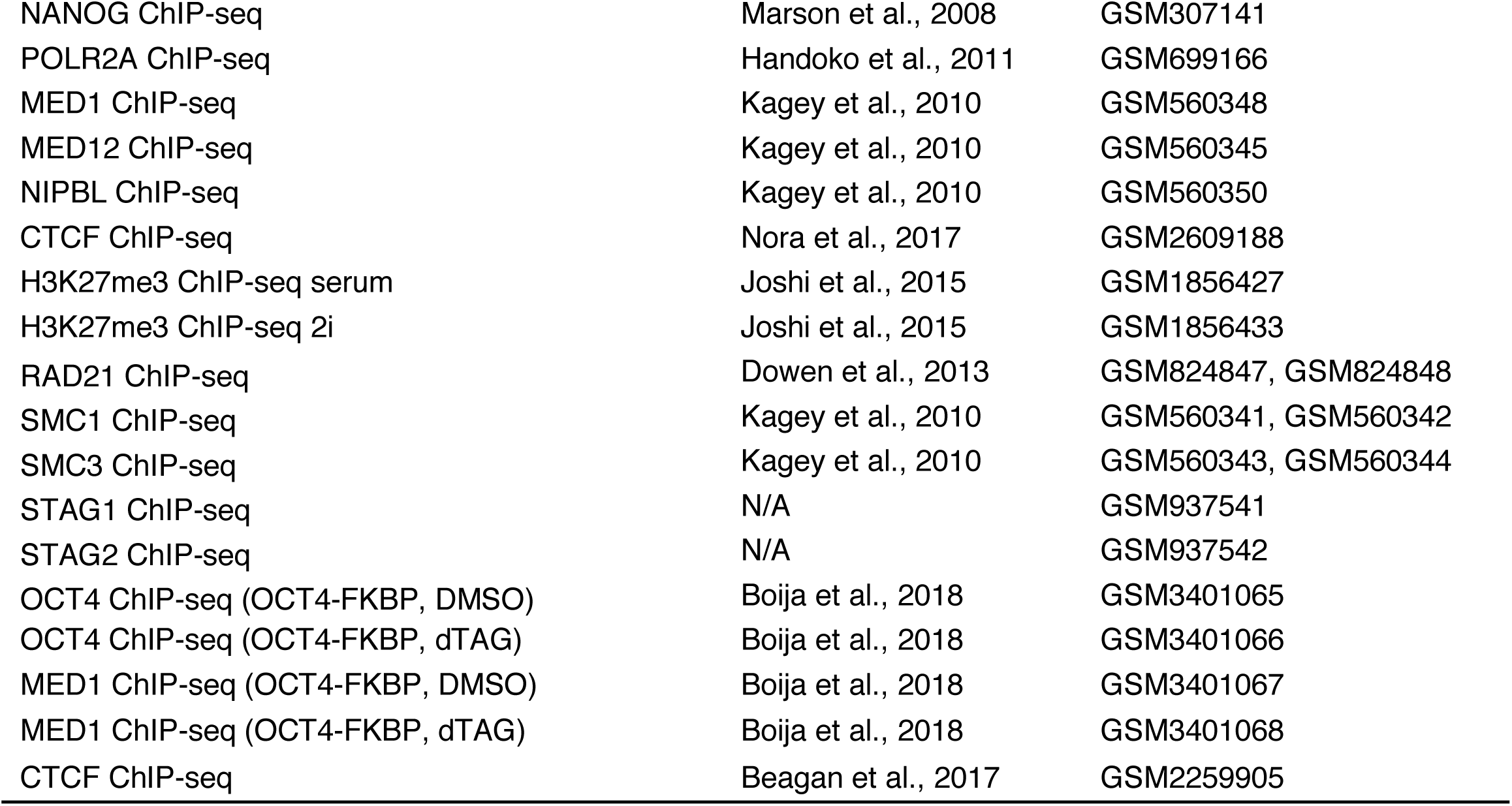

### Contact for Reagent and Resource Sharing

Further information and requests for reagents, plasmids and cell lines should be directed to the Lead Contact, Elzo de Wit.

### Experimental Model and Subject

#### Details Mouse Embryonic Stem Cells (ESCs)

E14Tg2a (129/Ola isogenic background) and the derived cell lines were cultured on 0.1% gelatin-coated plates in serum-free DMEM/F12 (Gibco) and Neurobasal (Gibco) medium (1:1) supplemented with N-2 (Gibco), B-27 (Gibco), BSA (0.05%, Gibco), 10^4^ U of Leukemia Inhibitory Factor/LIF (Millipore), MEK inhibitor PD0325901 (1 *µ*M, Selleckchem), GSK3-β inhibitor CHIR99021 (3 *µ*M, Cayman Chemical) and 1-Thioglycerol (1.5×10^-4^ M, Sigma-Aldrich). The cell lines were passaged every 2 days in daily culture. During the protein depletion experiments, the cells were seeded overnight before the start of the time course in the following densities: For a 96 h time course, 2.5 k, 35 k, 150 k, and 400 k cells were seeded in 24-well, 6-well, 10-cm and 15-cm plates, respectively. For 24 h time course, 5 k, 0.5 M, and 4 M cells were seeded in chamber slide (ThermoFisher Scientific), 6-well and 15-cm plates, respectively. The media were refreshed or the cells were split in 1:10 every 2 days during a time course.

#### Neural Progenitor Cells (NPCs)

The *OsTir1* parental and WAPL-AID cells were seeded at 100 k cells and cultured in serum-free medium without LIF and 2i. After 7 days, the cells were transferred on a 3.5-cm gelatinized (0.15% gelatin) plate and cultured in presence of recombinant murine EGF (10 ng/ml, PeproTech) and recombinant human FGF-basic (10 ng/ml, PeproTech) for an additional 7-10 days. The medium was refreshed daily during the differentiation procedure. The obtained neural progenitor cells were cultured on 0.1% gelatin-coated plates in the medium supplemented with EGF and FGF-basic and passaged every 3-4 days.

#### Indole-3-acetic Acid (IAA) and dTAG-13 Treatment

WAPL and RAD21 depletion were induced by treating the cells with a final concentration of 500 *µ*M IAA (I5148-10G, Sigma Aldrich). OCT4- and NANOG-FKBP proteins were depleted by adding a final concentration of 500 nM dTAG-13 molecule (requested from Dr. Nathanael S. Gray from Dana-Faber Cancer Institute) (Nabet et al., 2018). All the time series experiments were performed by inducing protein degradation at different time points and harvest the samples in the end of the time course.

### Method Details

#### Plasmid Construction

The donor plasmid used to target the endogenous mouse WAPL and NANOG protein was constructed by modifying a published pEN84 plasmid (Plasmid #86230, Addgene). Two homology arms around the stop codon of the Wapl genes were amplified by PCR from genome DNA of the *OsTir1* parental E14Tg2a cells. Two homology arms of the Nanog gene (3’ end) and FKBP^F36V^-HA-2A sequence were purchased from Integrated DNA Technologies, Eurofins Genomics and Twist Bioscience, respectively. To construct the *Wapl* donor plasmid, the AID-eGFP tag linked with a puromycin resistance gene driven by a PGK promoter (AID-eGFP-PuroR) and the backbone sequence were PCR amplified from the pEN84 vector. The homology arms, AID-eGFP-PuroR and the backbone were then assembled using Gibson Assembly Cloning Kit (E5510S, New England BioLabs), followed by replacing the PuroR into a Neomycin/Kanamycin resistance gene. Construction of the donor plasmid for *Rad21* targeting is similar to Wapl targeting with a replacement of *Wapl* homology arms and PuroR into *Rad21* homology arms and blasticidin resistance gene (*BlastR*), respectively. To construct the donor plasmid for Nanog targeting, the homology arms, the FKBP^F36V^-HA-2A sequence, the eGFP sequence and the backbone sequence were assembled using the same method as used for the *Wapl* donor plasmid.

To modify the Wapl gene, two sgRNAs were designed to target 3’-end sequence of the mouse Wapl gene. The Wapl-targeting sgRNAs were annealed the oligos caccgTCACTCTAGAGATAGACTTC and aaacGAAGTCTATCTCTAGAGTGAc and the oligo caccgTTACCTTTGCTTCAGGTGCT and aaacAGCACCTGAAGCAAAGGTAAc for the first and second sgRNA, respectively, and consequently cloned into a pX335 dual nickase plasmid (Plasmid #42335, Addgene). The sgRNA sequence CCACGGTTCCATATTATCTG was cloned into a pX330 plasmid (Plasmid #42230, Addgene) for *Rad21* target. To target the Nanog gene, a pair of annealed oligos, caccgTATGAGACTTACGCAACATC and aaacGATGTTGCGTAAGTCTCATAc, was cloned into a pX330 plasmid (Plasmid #42230, Addgene).

The donor sequences and sgRNAs in the obtained plasmids were validated by Sanger sequencing before using for further experiments.

#### Gene Targeting

The donor plasmids and their corresponding sgRNAs for *Wapl* and *Nanog* targeting were co-transfected into the parental cell lines using Lipofectamine 3000 Reagent (TheromFisher Scientific). Two to three days after transfection, the eGFP positive cells were sorted into a gelatinized 96-well plate for single clone selection. The obtained clones were genotyped by PCR and the fusion sequences were validated by Sanger sequencing. For *Rad21* targeting, the donor plasmid and the sgRNA were electroporated into wild-type E14Tg2a cells using Neon Transfection System (ThermoFisher Scientific). The transfected cells were selected with 10 *µ*g/ml blasticidin for 10 days, and then the *BlastR* was removed by transiently expressing flippase to trigger FRT recombination. Colonies were manually picked and genotyped by PCR for homozygous insertion of AID-GFP. An obtained homozygous RAD21-AID-eFP clone was electroporated in presence of 15 *µ*g of an *OsTIR1* donor plasmid (Plasmid #92142, Addgene) and 5 *µ*g of a sgRNA plasmid targeting endogenous *Tigre* locus. Clones were manually picked and grew in a 96-well plate, and further validated by PCR and flow cytometry.

#### Western Blots

mESCs and NPCs were harvested and lysed in RIPA lysis buffer (150 mM NaCl, 1% NP-40, 0.5% sodium deoxycholate, 0.1% SDS, and 25 mM Tris (pH=7.4)). The 6% in-house made SDS-PAGE gels were used to separate the Wapl and Rad21 proteins, and the 10% SDS-PAGE gels was used for Sox2, Oct4 and Nanog. The separated protein was transferred to a pre-activated PVDF membrane using Trans-Blot Turbo Transfer System (Bio-Rad). The blots were incubated with the following primary antibodies overnight at 4°C: (1) WAPL (1:1000, 16370-1-AP, Proteintech), (2) RAD21 (1:1000, ab154769, Abcam), (3) SOX2 (1:1000, D9B8N, Cell Signaling), (4) OCT4 (1:1000, D6C8T, Cell Signaling), (5) NANOG (1:1000, D2A3, Cell Signaling), and (6) HSP90 (1:2000, 13171-1-AP). After incubation, the blots were washed 3 times with TBS-0.1% Tween-20. The blots were then incubated with secondary antibody against rabbit IgG at room temperature for 1 h, following by 3-time TBS-0.1% Tween-20 washing. The proteins attached with antibodies were hybridized with Clarity Western ECL Substrate reagent (Bio-Rad) and visualized in a ChemiDoc MP Imaging System (Bio-Rad).

#### GFP Quantification and Cell Cycle Analysis

To quantify GFP signaling of Wapl depletion experiment, the WAPL-AID cells were treated with 500 nM IAA for 8 different time points (0, 5, 10, 20, 30, 45, 60, and 120 min), harvested and fixed with 2% paraformaldehyde at room temperature for 15 min. The parental cell line was also processed as a negative control. GFP signal was quantified on BD LSRFortessa analyzer (BD Biosciences).

Cell cycle analysis was performed following the protocol of Click-iT EdU Alexa Fluor 647 Flow Cytometry Assay Kit (Invitrogen). Briefly, cells were labeled with 10 *µ*M Click-iT EdU for 1.5 h. The cells were then fixed and permeabilized. The EdU was detected using Click-iT Plus reaction cocktail for 30 min at room temperature and protected from light, and DNA content of the cells was stained with DAPI. DAPI and EdU signals were quantified on BD LSRFortessa analyzer.

#### Immunofluorescence Staining

For GFP visualization cells were grown on poly-L-lysine (Sigma Aldrich) coated chamber slides (ThermoFisher Scientific), fixed in 4% formaldehyde (FA) and nuclei were counterstained with Hoechst 33342 (ThermoFisher Scientific).

For RAD21 immunofluorescence analysis in mESCs, we let single cells adhere for 30 min on poly-L-lysine coated slides. Next, pre-extraction of the non-chromatin-associated RAD21 fraction was performed by incubation with 0.1% Triton X-100 in PBS for 1 min followed by fixation with 4% FA. Staining was performed with rabbit-anti RAD21 (Abcam, ab154769, 1:200) followed by incubation with goat anti-rabbit Alexa Fluor 647 (Abcam, 1:250). Nuclei were counterstained with 4’,6-Diamidino-2-Phenylindole (DAPI) (ThermoFisher Scientific). For NPCs, cells were grown on poly-L-lysine coated coverslips fixed in 4% FA and stained with mouse anti-Nestin (BD biosciences, 611659, 1:200) and rabbit anti-GFAP (DAKO, Z033429-2, 1:100) antibodies, followed by incubation with goat anti-mouse Alexa Fluor 488 and goat anti-rabbit Alexa Fluor 568 antibodies (both ThermoFisher Scientific, 1:250). Nuclei were counterstained with DAPI. Prior to imaging all samples were mounted with FluorSave reagent (Merck). Fluorescent confocal images were captured on a Leica SP5 system (Leica, Wetzlar, Germany).

#### Alkaline Phosphatase Staining

Alkaline phosphatase staining was performed following the protocol of Leukocyte Alkaline Phosphatase Kit (Sigma-Aldrich). Cells were fixed in Citrate-Acetone-Formaldehyde solution for 30 s and gently washed in deionized water for 45 s, followed by stained in diluted Naphthol AS-BI Alkaline Solution at room temperature for 15 min, and visualized under bright-field microscopy.

#### ChIP-seq

All the ChIP-seq experiments, except for H3K27me3, were performed in presence of 10% HEK293T cells as an internal reference using a published protocol with small modifications (Liu et al., 2017). For chromatin preparation, the mouse embryonic stem cells were mixed with 10% HEK293T cells and cross-linked by a final concentration of 1% formaldehyde for 10 min. The cross-linking reaction was quenched using 2.0 M glycine. The cross-linked cells were then lysed and sonicated to obtain ∼300 bp chromatin using Bioruptor Plus sonication device (Diagenode). For ChIP assays, antibodies were first coupled with Protein G beads (ThermoFisher Scientific), and then the sonicated chromatin were incubated overnight at 4°C with the antibody coupled Protein G beads. After over incubation, captured chromatin was washed, eluted and de-crosslinked. The released DNA fragments were purified using MiniElute PCR Purification Kit (Qiagen). The ChIP experiments were performed using the following antibodies: (1) WAPL (16370-1-AP, Proteintech), (2) CTCF (07-729, Merck Millipore), (3) RAD21 (ab154769, Abcam), (4) SOX2 (AF2018, R&D Systems), (5) OCT4 (AF1759, R&D Systems), (6) NANOG (RCAB002P-F, Cosmo Bio Co.), (7) MED1 (A300-793A, Bethyl Laboratories), (8) H3K4me3 (pAb-003-050, Diagenode), (9) H3K27ac (ab4729, Abcam), and (10) H3K27me3 (pAb-195-050, Diagenode).

The purified DNA fragments were prepared according to the protocol of KAPA HTP Library Preparation Kit (Roche) prior to sequencing. All the ChIP-seq libraries were sequenced using the single-end 65-cycle mode on an Illumina HiSeq 2500.

#### RNA-seq

RNA was isolated following a standard TRIzol RNA isolation protocol (Ambion). The cells were lysed using 1 ml of TRIzol reagent, and 200 *µ*l chloroform was added to the lysates. The mixture was vortexed and centrifuged at 12,000 g at 4°C for 15 min. Upper phase was homogenized with 0.5 ml of 100% isopropanol, incubated at room temperature for 10 min, and centrifuged at 4°C for 10 min. The resulted RNA pellet was washed with 75% ice-cold ethanol, dried at room temperature for 10 min, and resuspended in RNase-free water. The isolated RNA was treated with DNase using RNeasy Mini Kit (Qiagen).

RNA-seq libraries were prepared using a TruSeq Stranded RNA LT Kit (Illumina). The libraries were sequenced using the same platform as the ChIP-seq libraries.

#### Hi-C

We generated Hi-C data as previously described (Rao et al., 2014) with minor modifications (Haarhuis et al., 2017). For each template, 10 million cells were harvested and crosslinked using 2% formaldehyde. Crosslinked DNA was digested in nucleus using MboI, and biotinylated nucleotides were incorporated at the restriction overhangs and joined by blunt-end ligation. The ligated DNA was enriched in a streptavidin pull-down. Hi-C libraries were prepared using a standard end-repair and A-tailing method and sequenced on an Illumina HiSeq X sequencer generating paired-end 150 bp reads.

#### 4C-seq

We generated 4C data for untreated and 24h IAA treated Wapl-AID and Rad21-AID cells. 4C was performed as previously described (Geeven et al., 2018; van de Werken et al., 2012) using a two-step PCR method for indexing described first in (Haarhuis et al., 2017). We used MboI as the first restriction enzyme and Csp6I as the second restriction fragment. Viewpoint specific primers can be found in the section of STAR&Methods. The 4C-seq libraries were sequenced using the same platform as the ChIP-seq libraries.

### Quantification and Statistical Analysis

#### ChIP-seq Analysis

Calibrated ChIP-seq data were analyzed based on a modified method described in previously (Orlando et al., 2014). Raw sequencing data was mapped to a concatenated reference genome (mm10 and hg19) using Bowtie 2 mapper (version 2.3.4.1) (Langmead et al., 2009). The mapped reads with mapping quality score <15 and the optical PCR duplicates were discarded using SAMtools (version 1.9) (Li et al., 2009). The reads derived from the reference HEK293T cells (hg19, raw human reads/HR^raw^) were scaled to 1 M reads which resulted in a scaling factor for normalizing the reads from mouse embryonic stem cells (mm10, raw mouse reads/MR^raw^). The scaling method can be summarized using the following steps:

1. derive a scaling factor (SF): SF = 1,000,000/HR^raw^;
2. compute scaled ChIP-seq coverage: MR^scale^ = MR^raw^ × SF, HR^scale^ = HR^raw^ × SF.

The coverage files (bigWig files) were generated by applying the above computed scaling factor using deepTools (version 2.5.4) (Ramírez et al., 2014). Peak calling was performed using MACS2 (version 2.1.1.20160309) (Liu, 2014) at a q-value cutoff of 0.01.

The scaled coverage files are not corrected for intensity bias caused by quality difference of the individual ChIP-seq profiles. Therefore, we computed average enrichment of the ChIP-seq experiments under direct comparison using their spike-in reference. The ratio between average enrichment of the spike-in reference was used to normalize the corresponding ChIP-seq profiles (see an example in Figure 1E).

#### Standard ChIP-seq Analysis

ChIP-seq data of H3K27me3 was performed without the presence of spike-in reference. The H3K27me3 ChIP-seq profiles and the re-analyzed publicly available ChIP-seq data were mapped to a mm10 reference. The mapped reads with mapping quality score <15 and the optical PCR duplicates were discarded using SAMtools. Peak calling was performed using MACS2 at a q-value cutoff of 0.01. The coverage files of uncalibrated ChIP-seq data and the data of pluripotency factors (due to absence of these factors in HEK293T cells) were generated using “normalize to 1X genome coverage” methods in deepTools.

#### ChIP-seq Peak Alignment and Functional Annotation

Alignment of ChIP-seq signal was performed using deepTools v3.0 (Ramírez et al., 2016). “Scale-regions” methods was applied to align the signal coverage from broad regions (RDC, RSC, super enhancers, and H3K27me3). Heatmaps were directly made using deepTools. Alignment plots were generally made with aligned matrices that were further processed in R.

The RDC and RSC were annotated using a web-version GREAT analysis tool (version 3.0.0) (McLean et al., 2010) against Mouse Genome Informatics (MGI) database (Bult et al., 2010) using a “basal plus extension” method to link ChIP-seq peaks to their gene targets.

#### Motif Analysis

A merged peak list was created from Rad21 ChIP-seq data of the control and treated NPCs. The read coverage under the peaks was determined using “peakstats.py” function in SolexaTools (version 2.1). The peaks with at least 10 reads in both replicates were kept for further analysis. DESeq2 (version 1.18.1) (Anders and Huber, 2010) was used to normalize the filtered coverage data between the samples based on their size factors. A Wald test in DESeq2 was used to detect differential peaks between the control and treated samples using a FDR cutoff of 0.01 and a fold change of 2.

We performed motif identification on the peaks higher in the untreated samples (0h enriched) and the unchanged peak set from the DESeq2 analysis using the GimmeMotifs (van Heeringen and Veenstra, 2011) using the non-redundant GimmeMotifs database (v3.0). Next we calculated for every motif the frequency in the 0 h enriched peak set and the constant peak set. We normalized the motif frequency by dividing the individual motif frequency by to total number of identified motifs (relative motif frequency). We calculated the log2-enrichment score by calculating the ratio of the 0 h enriched relative motif frequency dividing the constant relative motif frequency. The p-value was calculated using the Fisher exact test on the following 2×2 table: for every motif M, we determine the number of the 0 h enriched peaks with or without M and for the constant peaks with or without M.

#### RNA-seq Analysis

Raw RNA-seq data were mapped against mm10 reference genome using a TopHat2 pipeline (version 2.1.1) (Kim et al., 2013). The mapped reads with mapping quality score <10 were discarded using SAMtools. The read coverage for each gene in “Mus Musculus GRCm38.92” annotation file was determined using a HTSeq tool (version 0.9.1). The coverage files were generated using “normalize to 1X genome coverage” methods in deepTools.

The genes with at least 20 reads in both replicates were kept for further analysis. The filtered expression data were normalized based on the size factors of the individual samples using a DESeq2 package. The significant genes were detected by comparing the control and treated samples using Wald test built in DESeq2 with an FDR of 0.05. The results were visualized using “heatmap.2” function in a “gplots” package (version 3.0.1.1). GSEA was performed using a desktop version of GSEA tool (version 3.0) (Subramanian et al., 2005) and a Molecular Signatures Database (MSigDB, version 6.2) (Liberzon et al., 2015). The genes were ranked based on the difference in log2 ratios between the control and treated samples. The seed for permutation was set at the option “149”.

#### Hi-C data Processing

Raw Hi-C data were mapped with HiC-Pro (Servant et al., 2015), which performs mapping, identification of valid Hi-C pairs, generation of contact matrices and ICE normalization (Imakaev et al., 2012). Subsequent analyses were performed in GENOVA, a Hi-C visualization tool written in R (http://github.com/deWitLab/GENOVA).

#### Self-interaction Score

In order to calculate the degree of local self-interaction we calculated a self-interaction score (*SI*). We used 20kb Hi-C matrices to calculate the *SI*. In order to calculate the degree of self-interaction for a given region *i* of window size *w*, we calculate the mean contact frequency of all the Hi-C bins in this region with each other. Effectively, this means calculating the average signal within a triangle close to the diagonal (Figure 4A). Note that the bottom of the triangle is nearest to the diagonal. Because the highest signal is on the diagonal itself we remove the diagonal from the self-interaction score. To correct for chromosome-wide trends in the self-interaction score, we subtract for every region *i* in the genome the median of the 100 self-interaction scores up and down from that window and the self-interaction score of that window itself (*SI*_*i*_ – median {*SI*_*i-100*_, *SI*_*i-99*_,…,*SI*_*i+100*_}). In this way the self-interaction score is calculated over a local background, explaining negative *SI* scores.

#### RDC/RSC RNA-seq Intersection

We intersected the RDCs and RSCs with the expression data by determining for every RDC and RSC the closest gene, from here on called RDC or RSC gene. Next, we determined for the RDC and RSC genes whether they are upregulated, downregulated or unchanged. The fraction of genes in every category (observed) was compared to the genome-wide fraction of genes in the upregulated, downregulated or unchanged category (expected). The ratio of observed over expected was calculated for every time point, RDC, RSC and cell line. To determine the probability of this happening by accident we performed a circular permutation analysis using regioneR (Gel et al., 2016). The confidence intervals and empirical p-values are the result of 10,000 permutations.

#### Identification of RDCs and RSCs

In order to identify RDCs and RSCs we binned the RAD21 ChIP-seq signal in untreated and 24 hour treated WAPL-AID cells to 100bp bins. Next we perform per chromosome quantile normalization(Bolstad et al., 2003). We calculate the difference between untreated and the treated and discretize into three observation values: ‘ChIP_up’ (difference between 0h and 24h > 1), ‘ChIP_down’ (difference between 0h and 24h < −1) and ‘ChIP_same’ (difference between 0h and 24h > −1 and < 1). We create a fully connected hidden markov model with three states: RDC, RSC and no_change. Every state has specific emission probabilities for the different observations and transition probabilities of 10^-6^ to transition into a different state. This analysis is implemented using the function from the R package HMM.

#### 4C-seq Analysis

The raw sequence data was mapped using our 4C mapping pipeline (http://github.com/deWitLab/4C_mapping). We normalized our 4C data to 1 million intrachromosomal reads and visualize chromatin interactions around the viewpoints using peakC (http://github.com/deWitLab/peakC) (Geeven et al., 2018).

## Figure legends

**Supplementary Figure 1: Generating a WAPL-AID degron line.** A) Homologous recombination strategy for tagging the endogenous *Wapl* gene with AID-eGFP. Note that the targeting construct contains a *NeoR/KanR* resistance gene, but that this has not been used for selecting the first clone. Middle panel shows PCR validation (primers highlighted in left panel) showing homozygous integration of the donor vector. Right panel shows Sanger sequencing results confirming in-frame tagging of AID tag. B) Example loci showing WAPL ChIP-seq in WAPL-AID mouse ESCs and human HEK-293T cells following WAPL depletion in WAPL-AID cells.. C) Western blot showing the protein levels of the pluripotency transcription factors OCT4, SOX2 and NANOG. D) FACS plots showing DAPI staining and EdU incorporation to measure the cycle phase of individual cells. E) Quantification of cycle phases based on DAPI/EdU FACS analysis for various times after WAPL depletion. Error bars show standard deviation for triplicate experiments. F) Cell cycle profile as measured by DAPI staining for WAPL depletion time series. G) Morphology analysis and alkaline phosphatase staining for a second WAPL-AID clone similar to Figure 1F,G.

**Supplementary Figure 2: WAPL depletion does not cause Polycomb-associated epigenetic changes.** A) MA-plot showing average expression versus differential expression for various timepoints. Genes showing differential expression (FDR < 0.05) are highlighted in red (upregulation) or blue (downregulation). B) Heatmap showing the expression of well-studied pluripotency genes. Right panel shows the quantification of the differential expression. C) GSEA similar to Figure 2B but for the Chemical and Genetic Perturbation geneset from the MSigDB database. The top 10 ranked by -log_10_ p-value is highlighted in detail. D) ChIP-seq profiles for H3K27me3 peaks identified in serum, representing the canonical PRC2 target regions. First two columns show published H3K27me3 in serum and 2i conditions. Last two columns show data for H3K27me3 levels in treated and untreated WAPL-AID cells (both grown in 2i). E) HoxD locus shown as an example for H3K27me3 levels for the datasets shown in D).

**Supplementary Figure 3: Further characterization of the Regions of Dynamic Cohesin.** A) Venn diagram showing the uniquely bound and overlapping RAD21 binding sites identified by ChIP-seq in a second WAPL-AID clone. Peak calling was performed on a subset of 7 M sequencing reads for both of the samples. B) Barplot showing the number of RDCs and RSCs identified by the hidden markov model analysis. C) Alignment of RAD21 binding data from a second WAPL-AID clone on the RDCs and RSCs identified in the original WAPL-AID clone. Alignment of published ChIP-seq data for five different cohesin subunits generated in V6.5 mESC line (see STAR Methods for details). E) Example locus showing a region with an RDC and RSC. RAD21 binding is shown for treated and untreated cell for two WAPL-AID clones. Cohesin binding is shown for the five subunits shown in D). F) RAD21 binding as measured by ChIP-seq aligned to super/stretch enhancers for treated and untreated cells in two different clones. SE positions were taken from dbSUPER (Khan and Zhang, 2016). G) RAD21 alignment on the transcription start sites (TSS) of the top 1000 most stably expressed housekeeping genes (de Jonge et al., 2007) for treated and untreated cells in two different clones.

**Supplementary Figure 4: WAPL depletion reorganizes the 3D genome.** A) Relative contact probability plot (RCP) showing the distribution of the chance of two loci contacting in the context of the 3D genome as a function of the distance between them on the linear chromosome. B) Example locus showing the formation of extended loops upon WAPL depletion. Heatmaps visualize the contact frequency matrices for the *Nfe2l3* locus. Rightmost panel shows the differential contacts between the 24h treated cells and the untreated cells. C) Example regions showing a drop in contact frequency for the RDCs in the *Klf9* locus. D) High-resolution 4C data showing the contact profile of the *Klf9* promoter for the untreated and treated cells, with the differential contact profile below. ChIP-seq tracks show the binding of RAD21 in two independent clones for this region. E) Insulation score alignments for RDCs and RSCs in untreated and treated cells. For definition of insulation score see (Crane et al., 2015). F) Example plot showing the binding distribution of RAD21 in untreated and untreated cells and the insulation scores for the same region. Insulation scores have been plotted for a range of window sizes visualized in a domainogram style.

**Supplementary Figure 5: Effects of WAPL and RAD21 depletion on expression, transcription factor binding and genome organization.** A) Observed over expected ratios for genes differentially expressed genes or unchanged in WAPL-AID cells nearest to an RSC. B) Alignment of ChIP-seq data for pluripotency transcription factors SOX2 and OCT4 to RDCs and RSCs in untreated and treated cells to RDCs and RSCs. C) High-resolution 4C analysis for the *Elf3* locus in WAPL-AID cells. 4C data plotted in the same manner as in Figure 4F. ChIP-seq track show binding of NANOG and MED1 in untreated and treated cells. RNA tracks show the expression of *Elf3* in treated and untreated cells. D) Western blot showing acute depletion of RAD21. E) Bright-field microscopy images showing the morphology of RAD21-AID cells before and after 24 hours of RAD21 depletion. F) Western blot showing the levels of pluripotency transcription factors OCT4, SOX2 and NANOG. G) Violin plot similar to Figure S2B quantifies the expression of pluripotency factors in RAD21 depletion experiments. H) Observed over expected ratios for genes differentially expressed genes or unchanged in RAD21-AID cells nearest to an RSC. I) Alignment of NANOG and MED1 ChIP-seq data on RDCs and RSCs in treated and untreated RAD21-AID cells. J) High-resolution 4C data plotted similar to C) but for RAD21-AID cells.

**Supplementary Figure 6: The role of transcription factors in the recruitment of cohesin.** A) ChIP-seq heatmaps showing RAD21 and MED1 binding in the OCT4-FKBP line. RAD21 levels are plotted for OCT4 binding sites defined by (Boija et al., 2018). B) ChIP-seq heatmaps showing RAD21 binding at CTCF binding sites in OCT4-FKBP and NANOG-FKBP. For OCT4-FKBP CTCF sites from V6.5 mESCs (Beagan et al., 2017) were used for alignment and for Nanog-FKBP, which are derived from E14 mESCs CTCF sites identified in WAPL-AID cells were used. C) MED1 ChIP-seq alignment over RDCs and RSCs following OCT4 depletion. D) RAD21 and MED1 ChIP-seq alignment over RDCs and RSCs following NANOG depletion. E) The binding of RAD21 and MED1 over an RDC in treated and untreated OCT4-FKBP and NANOG-FKBP cells is shown for the *Esrrb* locus. OCT4 ChIP-seq track shows previously published OCT4 profiles in the same cell line (Boija et al., 2018).

**Supplementary Figure 7: RAD21 binding in WAPL-AID NPCs.** A) Staining for neuronal markers in NPCs derived from the parental *OsTir1* cell line. B) Staining for alkaline phosphatase in NPCs derived from mESCs. C) Venn diagram shows overlap of RAD21 binding sites measured ChIP-seq for untreated and 24h treated WAPL-AID NPCs. Only sites that showed consistent unique or shared binding across two replicates were included. Peak calling was performed on a subset of 7 M sequencing reads for all the samples. D) MA-plot showing the log_2_ fold-change as a function of the number of reads in a RAD21 peak. Highlighted dots indicate significantly changed RAD21 levels based on DESeq2 with an FDR <0.01 and a fold-change of at least 2 up or down. Red is increase in RAD21, blue is decrease.

