## Supplementary figures and images for "WAPL maintains dynamic cohesin to preserve lineage specific distal gene regulation"

### Supplemental Figures

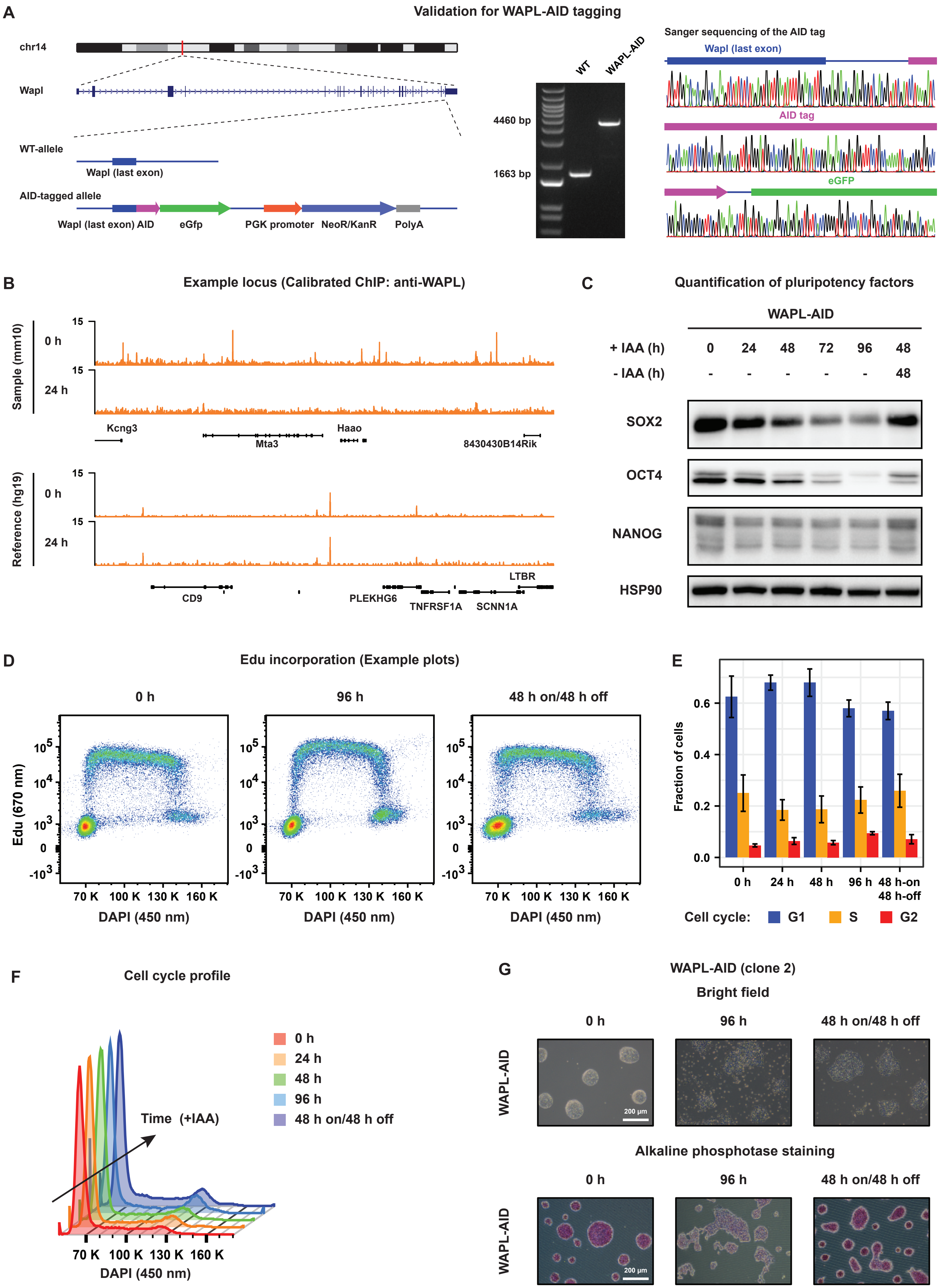

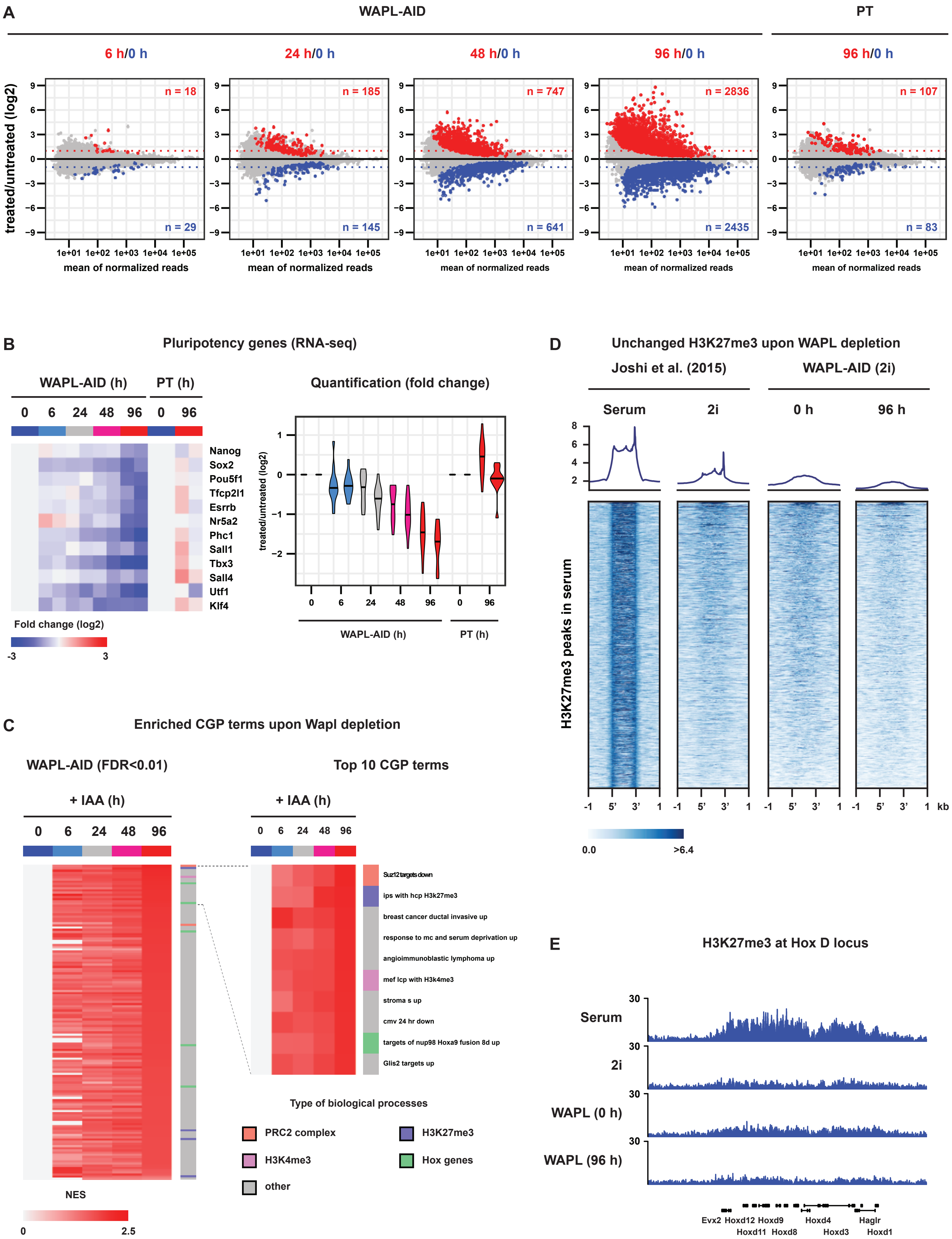

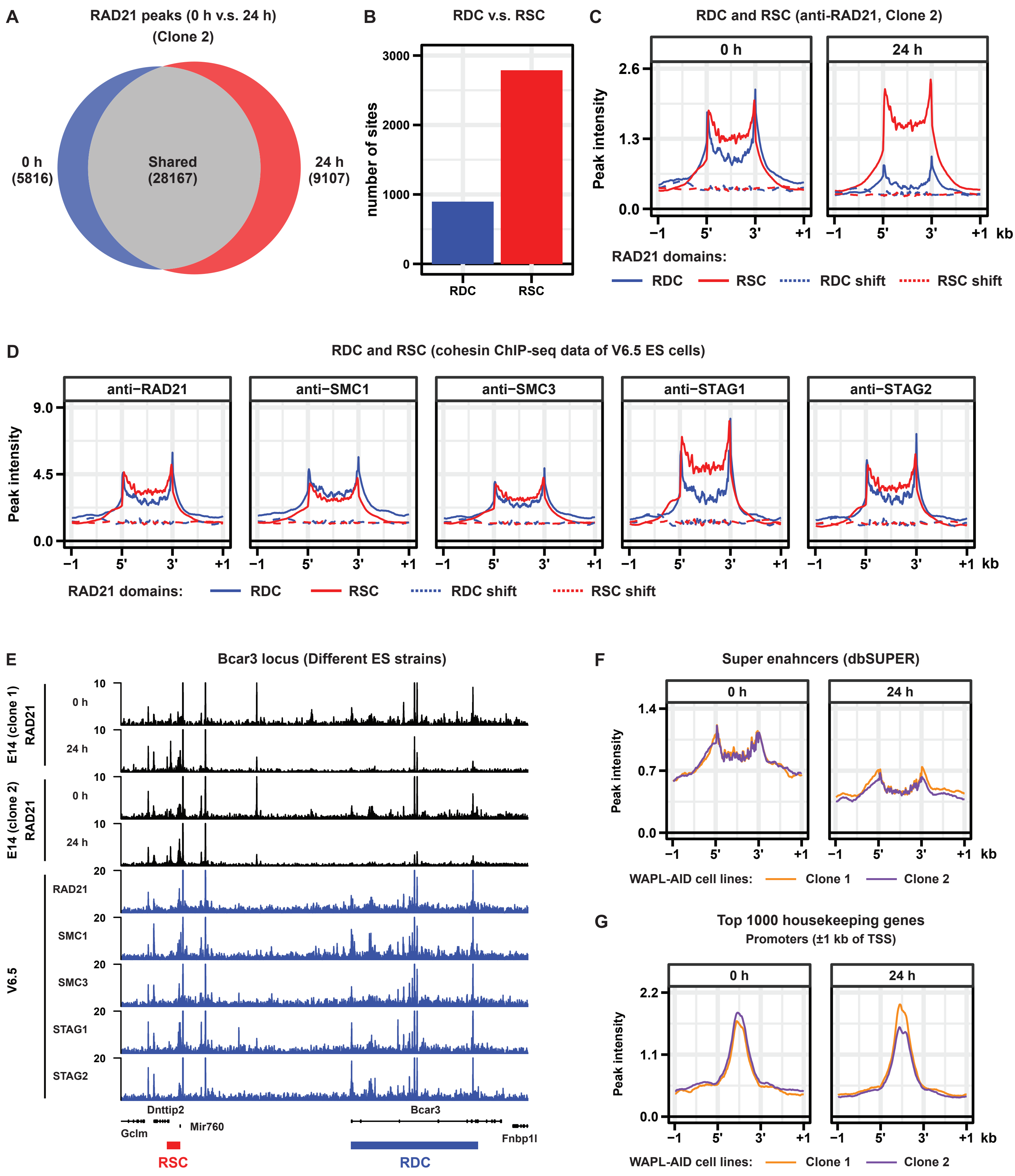

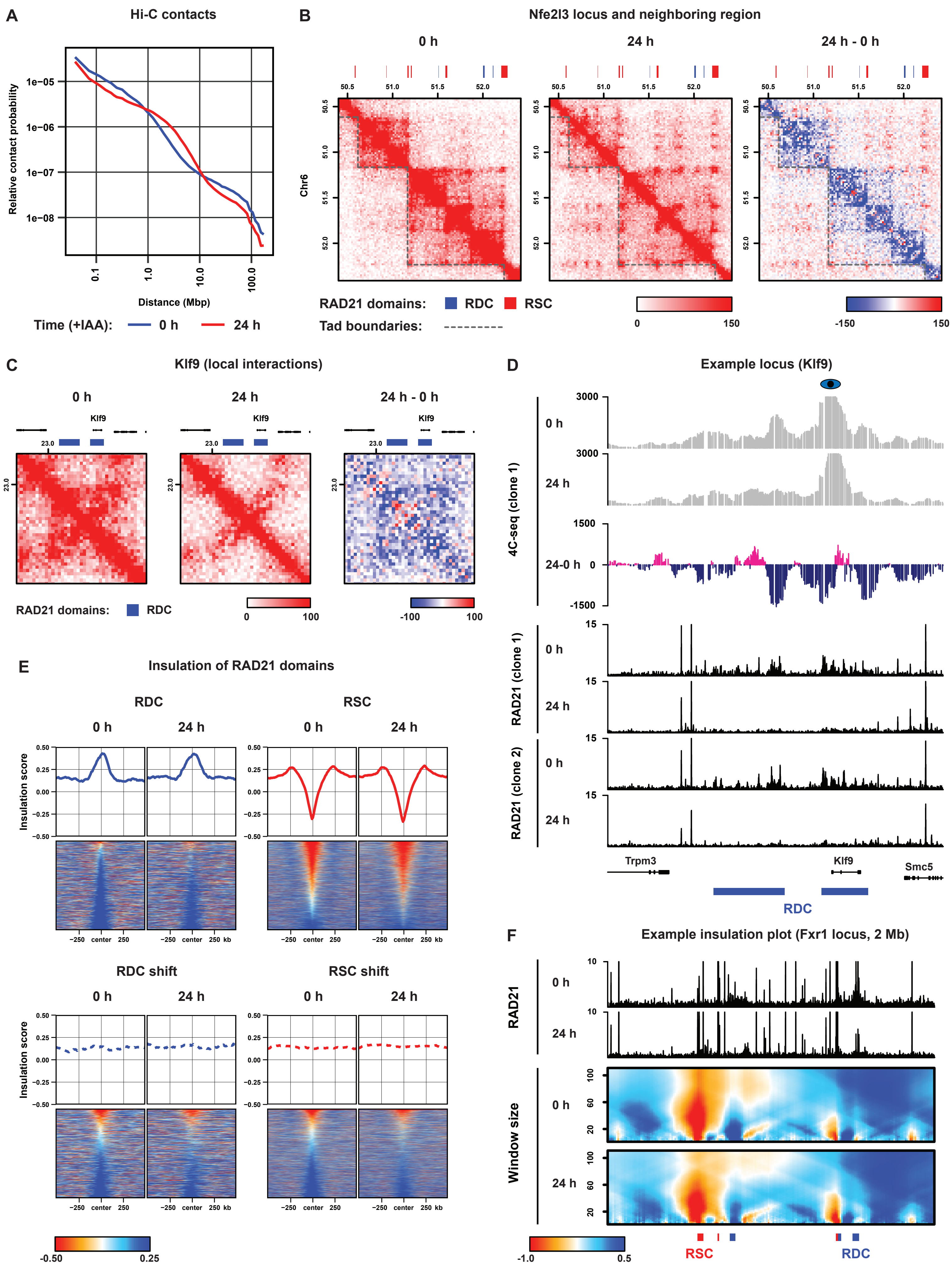

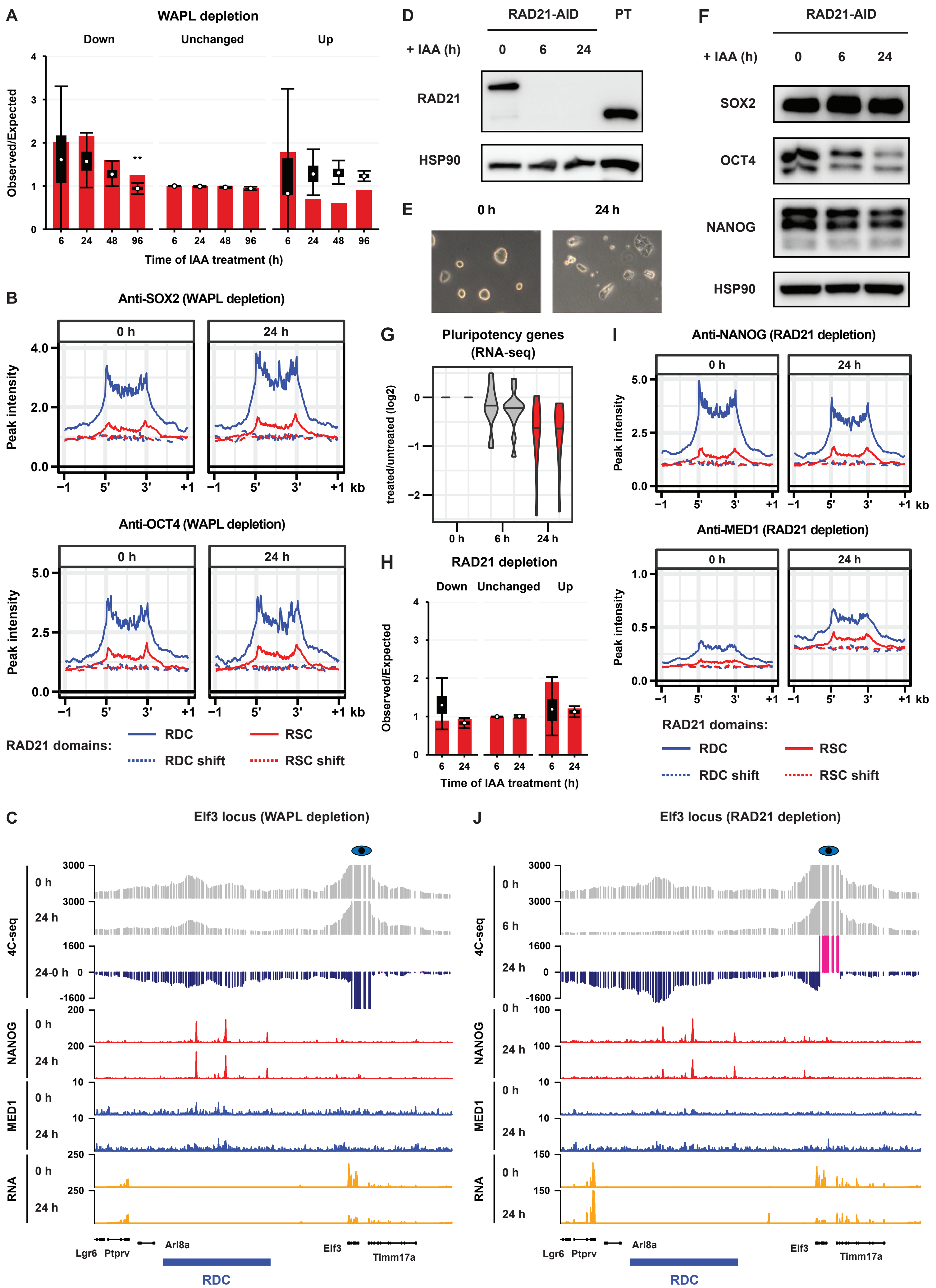

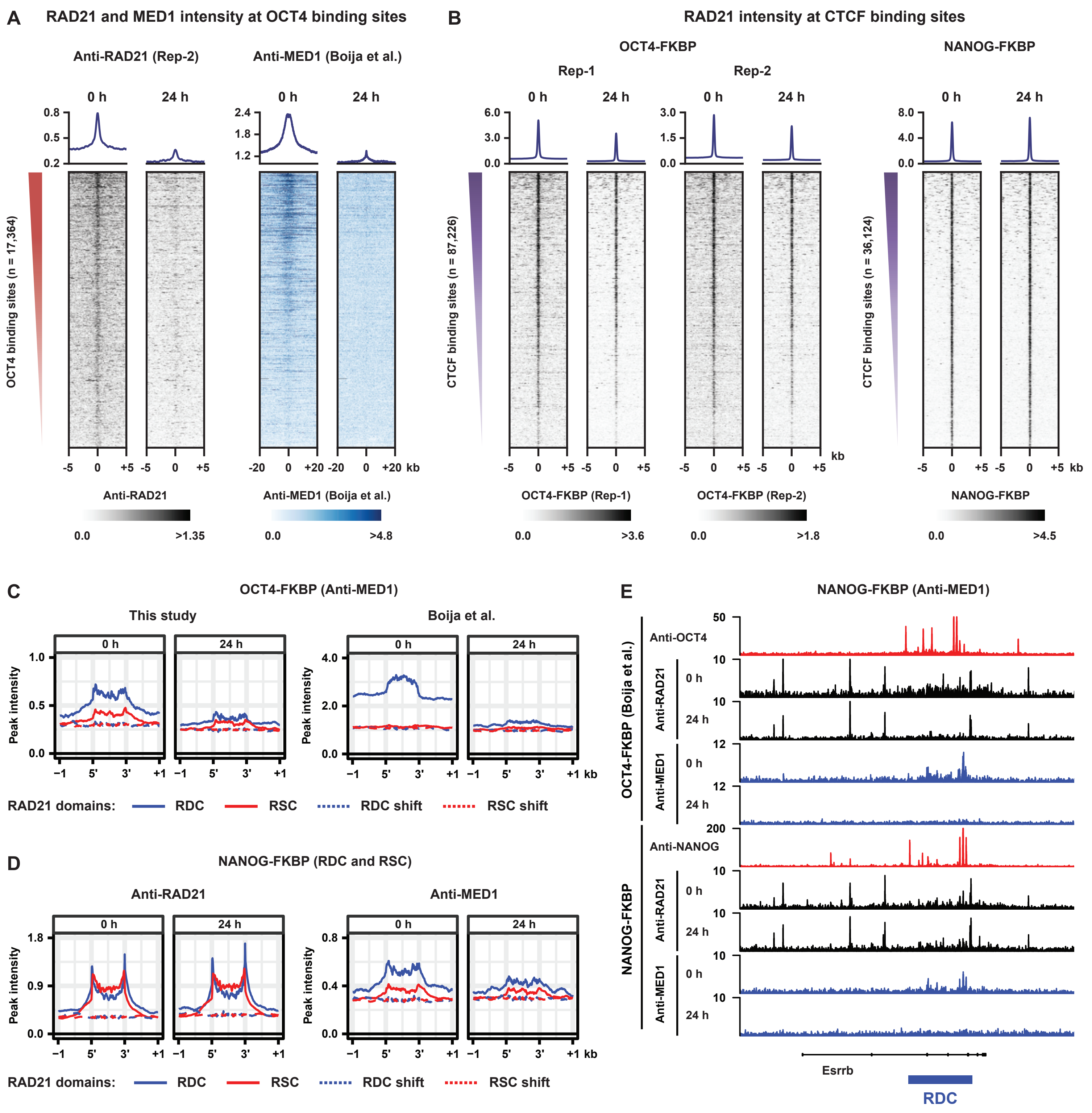

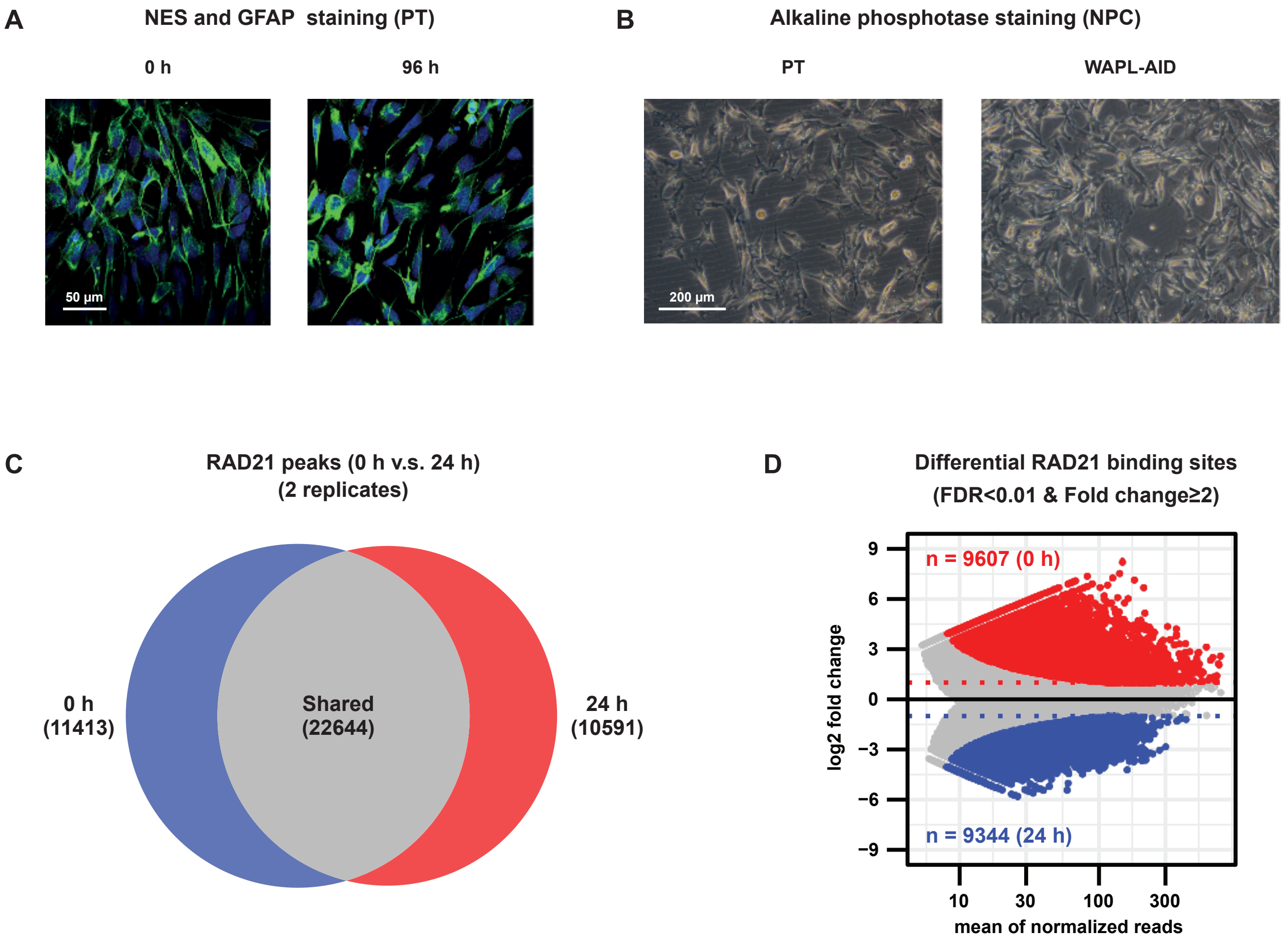
